# Linking Cerebellar Functional Gradients to Transdiagnostic Behavioral Dimensions of Psychopathology

**DOI:** 10.1101/2020.06.15.153254

**Authors:** Debo Dong, Xavier Guell, Sarah Genon, Yulin Wang, Ji Chen, Simon B. Eickhoff, Cheng Luo, Dezhong Yao

**Author notes:** Corresponding to: School of life science and technology, University of Electronic Science and Technology of China, Chengdu, China; (C. Luo).

## Abstract

High co-morbidity and substantial overlap across psychiatric disorders encourage a transition in psychiatry research from categorical to dimensional approaches that integrate neuroscience and psychopathology. Cerebellum is involved in a wide range of nonmotor cognitive functions and mental disorders. An important question thus centers on the extent to which cerebellar function can be linked to transdiagnostic dimensions of psychopathology. Here, this question is investigated using partial least squares to identify latent dimensions linking cerebellar connectome properties as assessed by macroscale spatial gradients of connectivity to a large set of clinical and behavioral measures in 198 participants across diagnostic categories. This analysis reveals significant correlated patterns of cerebellar connectivity gradients and behavioral measures that could be represented into four latent dimensions: general psychopathology, general lack of attention regulation, internalizing symptoms, and dysfunctional memory. Each dimension is associated with a distinct spatial pattern of cerebellar connectivity gradients. These findings highlight the relevance of cerebellar connectivity as a necessity for the study and classification of transdiagnostic dimensions of psychopathology.

## Introduction

Our understanding of cerebellar contributions to neurological function has changed from a traditional view focused on motor coordination, to a modern understanding that also implicates the cerebellum in a broad range of high-level cognitive and affective processes.^1^ An increasing body of evidence also supports cerebellar involvement in a wide range of psychiatric disorders.^2, 3^ Up to now, most psychiatric studies investigating the role of the cerebellum have been conducted based on categorical diagnostic criteria that view psychiatric disorders as independent entities.^4^ It is increasingly recognized that existing clinical diagnostic categories might be suboptimal, as there is substantial overlap in symptoms, cognitive dysfunction and genetic factors across multiple psychiatric disorders.^4, 5^ These overlaps can be reflected by shared neurobiological structure and function, and polymorphism abnormalities across psychiatric syndromes.^6–9^ The high rates of comorbidity between psychiatric disorders and heterogeneity within one diagnostic group further exacerbates this problem.^10–12^ This context has motivated transdiagnostic initiatives, such as the National Institute of Mental Health’s Research Domain Criteria,^13^ which encourages a transition in psychiatry research from categorical to dimensional approaches that integrate neuroscience and psychopathology.^13^

Recent clinical neuroscience studies have begun to adopt transdiagnostic approaches to highlight the importance of altered cerebellar structure in broad risk for psychopathology.^14–16^ Previous animal and human neuroimaging studies have provided converging evidence for the involvement of cerebellar function in a wide range of behaviors that are dependent on circuits connecting the cerebellum with multiple cerebral cortical regions.^1, 17–19^ Accumulating evidence supports dysfunctional cerebellar connectivity in many psychiatric disorders, such as schizophrenia,^20^ bipolar disorder,^21^ major depression,^22^ attention-deficit/hyperactivity disorder^23^ and autism.^24^ Moreover, study of clinical high-risk subjects demonstrate that dysconnectivity of cerebellar circuits can serve as a state-independent neural signature for psychosis prediction and characterization.^25^ Within this context, an understudied area of investigation is the extent to which cerebellar function can be linked to transdiagnostic dimensions of psychopathology.

Resting-state functional connectivity has been widely used to characterize disconnection mechanisms in many psychiatric disorders,^26, 27^ and is a promising tool for deepening our understanding of transdiagnostic dimensions.^28–30^ However, previous studies investigating functional connectivity-informed dimensions of psychopathology often ignore the importance of the cerebellum, e.g., by using a coarse delineation of the cerebellum with only a few regions of interest to represent the whole cerebellar information.^29, 30^ Recent developments in cerebellar functional mapping indicate that cerebellar functional organization can be characterized using macroscale spatial gradients of connectivity, a low dimensional continuous space that reflects the overarching spatial patterns that underpin the observed neural data.^31^ The principal connectivity gradient of cerebellar cortex captures a progression from sensorimotor to cognitive processing areas,^31^ similar to the organization of the cerebral cortex.^32, 33^ This low-dimensional representation of the principal axis of cerebellar macroscale functional organization thus provides a useful tool to characterize cerebellar function at the single-subject level which can then be correlated with single-subject behavioral measures. This approach offers an unprecedented opportunity to interrogate the relationship between cerebellar functional organization and behavioral measures of clinical phenomena, cognitive ability, and personality traits in mental health and disease.

In this study, we analyzed UCLA Consortium for Neuropsychiatric Phenomics open access dataset, a large resting-state fMRI and behavioral dataset^34^ using gradient-based and partial least squares, a multivariate data-driven statistical techniques with the objective to discover the latent dimensions that link cerebellar functional organization to behavioral measures spanning clinical, cognitive, and personality trait domains (Table S1 and Table S2) across healthy controls (HC, n=92) and patients with attention-deficit/hyperactivity disorder (ADHD, n=35), bipolar disorder (BD, n=36) and schizophrenia (SZ, n=35). Table 1 shows a summary of demographic and clinical information of each group. This approach yielded dimensions that optimally linked co-varying cerebellar connectivity gradients and behavior in individuals across traditional diagnostic categories, in accordance with a transdiagnostic dimensional approach. Multiple control analyses were used to optimize the robustness of these latent dimensions. Furthermore, we performed 10-fold cross-validation to assess the generalization performance of latent dimensions to unseen test data. Importantly, cross-validation approaches can help guard against overfitting that arises from high dimensional neurobiological data.^35^

**Table 1.** Demographic characteristics of 1 the each diagnostic group

| Variables | ADHD | BD | HC | SZ | F or $X^2$ | P value |
| --- | --- | --- | --- | --- | --- | --- |
| Sample size | 35 | 36 | 92 | 35 |  |  |
| Age (years, mean(SD)) | 31.40(10.50) | 34.44(8.91) | 30.50(8.50) | 35.54(8.97) | 3.51 | $1.6 \times 10^{-2}$ |
| Male sex, n(%) | 18(51.4) | 19(52.8) | 51(55.4) | 27(77.1) | 6.54 | $8.8 \times 10^{-2}$ |
| Education (years, mean(SD)) | 14.43(1.79) | 14.64(1.94) | 15.26(1.62) | 12.71(1.64) | 18.75 | $1.0 \times 10^{-10}$ |
| Site 1, n(%) | 17(48.6) | 18(50) | 73(79.3) | 14(40) | 23.72 | $2.9 \times 10^{-5}$ |
| Head motion, mean FD, mean(SD) | 0.069(0.04) | 0.083(0.05) | 0.066(0.03) | 0.096(0.04) | 6.16 | $5.1 \times 10^{-4}$ |
| Number of current medication use (mean(SD)) | 0.57(1.14) | 2.50(1.93) | 0(0) | 2.20(1.57) | 57.19 | $1.6 \times 10^{-26}$ |
| Number of substance use (mean(SD)) | 1.31(1.68) | 2.58(2.09) | 0.62(1.10) | 2.46(2.23) | 17.89 | $2.7 \times 10^{-10}$ |

Notes: Group differences were determined by either one-way ANOVA for continuous variables or chi-square tests for categorical variables. FD, framewise displacement; Number of substances use, including nicotine, alcohol, cannabis and other psychotropic substances

## Results

### Pattern of the principal functional connectivity gradient in cerebellum

The principal gradient (or principal gradient) explains as much of the variance in the data as possible (∼30%, Figure 1), represents a well-understood motor-to-supramodal organizational principle in the cerebellar connectivity. The principal connectivity gradient of cerebellar cortex captureed a progression from sensorimotor to cognitive processing areas. Specifically, it extended bilaterally from lobules IV/V/VI and lobule VIII to posterior aspects of Crus I and Crus II as well as medial regions of lobule IX. This observed spatial distribution was similar to previous reports of the principal functional conectivity gradient of the cerebellar cortex in healthy humans.^31^

**Figure 1.**
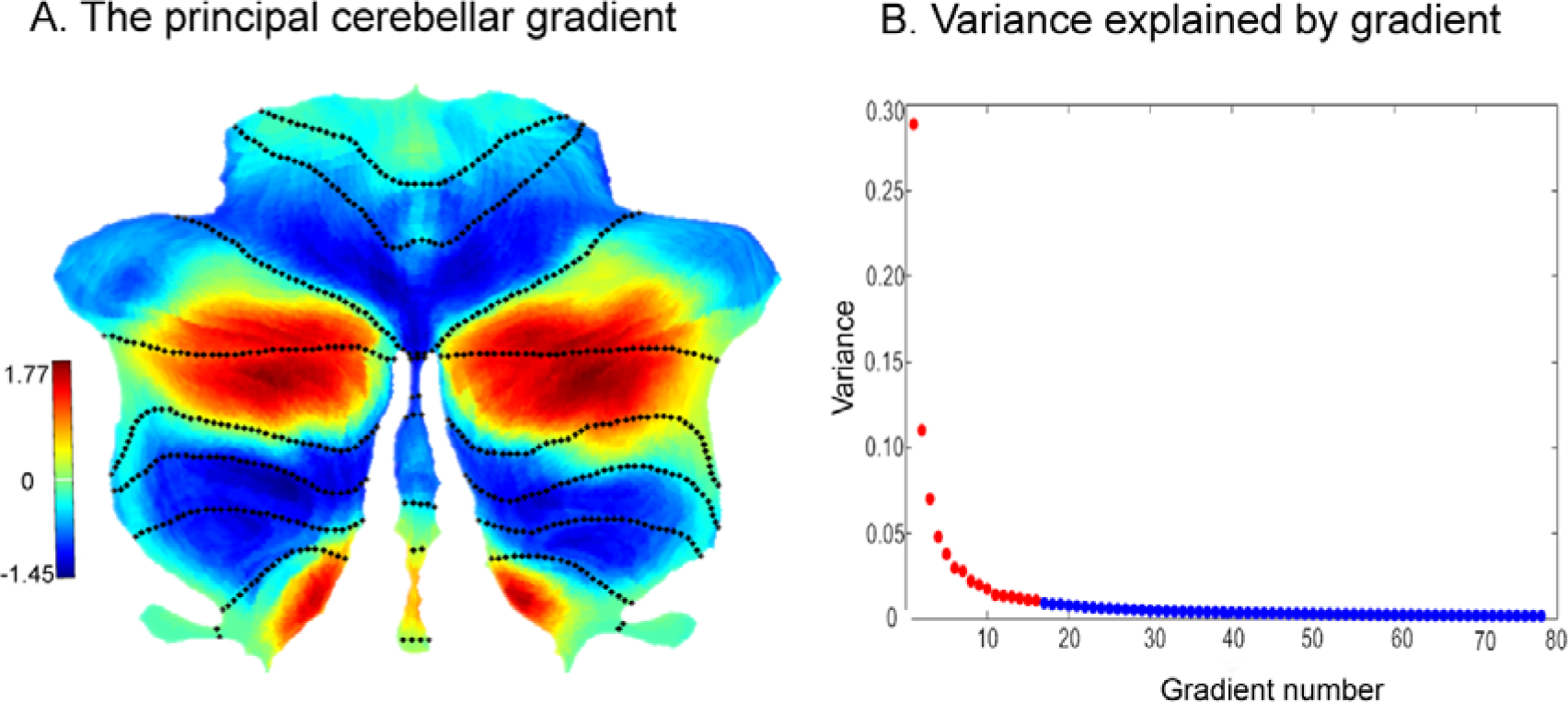
(A) The principal cerebellar connectivity gradient. (B) Covariance explained by each gradient. Red circles correspond to the gradients that explained at least a variance of 1%. Four Robust LVs Linking Cerebellar Gradients and Behavior

PLS correlation analysis revealed five significant latent variables (LVs) that reflect the direct covariant mapping between cerebellar connectivity gradients and behavioral measures. Since the fifth LV did not show robustness in control analyses as detailed in Table S3, we only focused on the first four LVs (LV1: r=0.62, permuted p=2.0×10^-2^; LV2: r=0.56, permuted p=2.0×10^-3^; LV3: r=0.61, permuted p=3.0×10^-2^; LV4: r=0.60, permuted p=1.2×10^-2^; **Figures 2, 3, 4, 5**A). The variance explained by each LV was 19.5%, 13.7%, 8.8% and 6.0%, respectively (Figure S1). Importantly, 10-fold cross-validation confirmed generalizability (i.e. robustness of results in new data) of the first four LVs, as indicated by significant correlation between cerebellar gradient and behavioral composite scores in the test folds (LV1, r=0.21, p=2.5×10^-3^; LV2, r=0.27, p=2.1×10^-3^; LV3, r=0.22, p=2.3×10^-3^; LV4, r=0.16, p=2.5×10^-3^).

**Figure 2.**
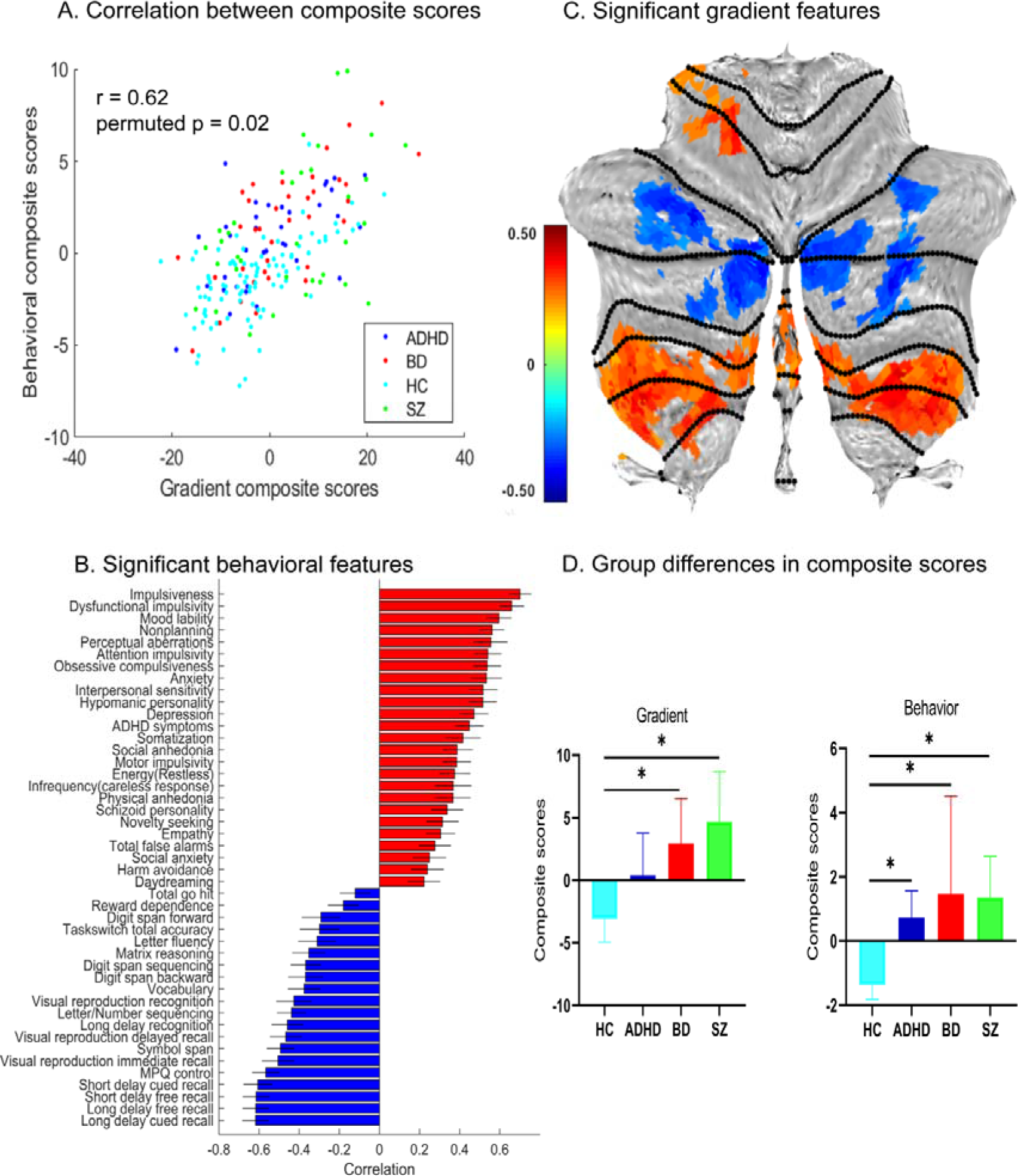
Latent variable 1: general psychopathology. (A) Correlation between cerebellar connectivity gradient and behavioral composite scores of participants. (B) Significant behavioral features associated with LV1. The contribution of each behavior is measured by correlations between participants’ behavioral scores and the corresponding behavioral composite scores. Error bars indicate bootstrapped standard deviations. (C) Significant gradient pattern associated with LV1. The contribution of each voxel is measured by correlation between participants’ cerebellar gradient scores and the corresponding cerebellar gradient composite scores (FDR correction, q<0.05). Gradient pattern displayed on cerebellar flat maps were generated using the SUIT toolbox (http://www.diedrichsenlab.org/imaging/suit.htm). (D) Group differences in cerebellar connectivity gradient and behavioral composite scores. Significant differences are indicated by asterisks (FDR correction, q < 0.05).

**Figure 3.**
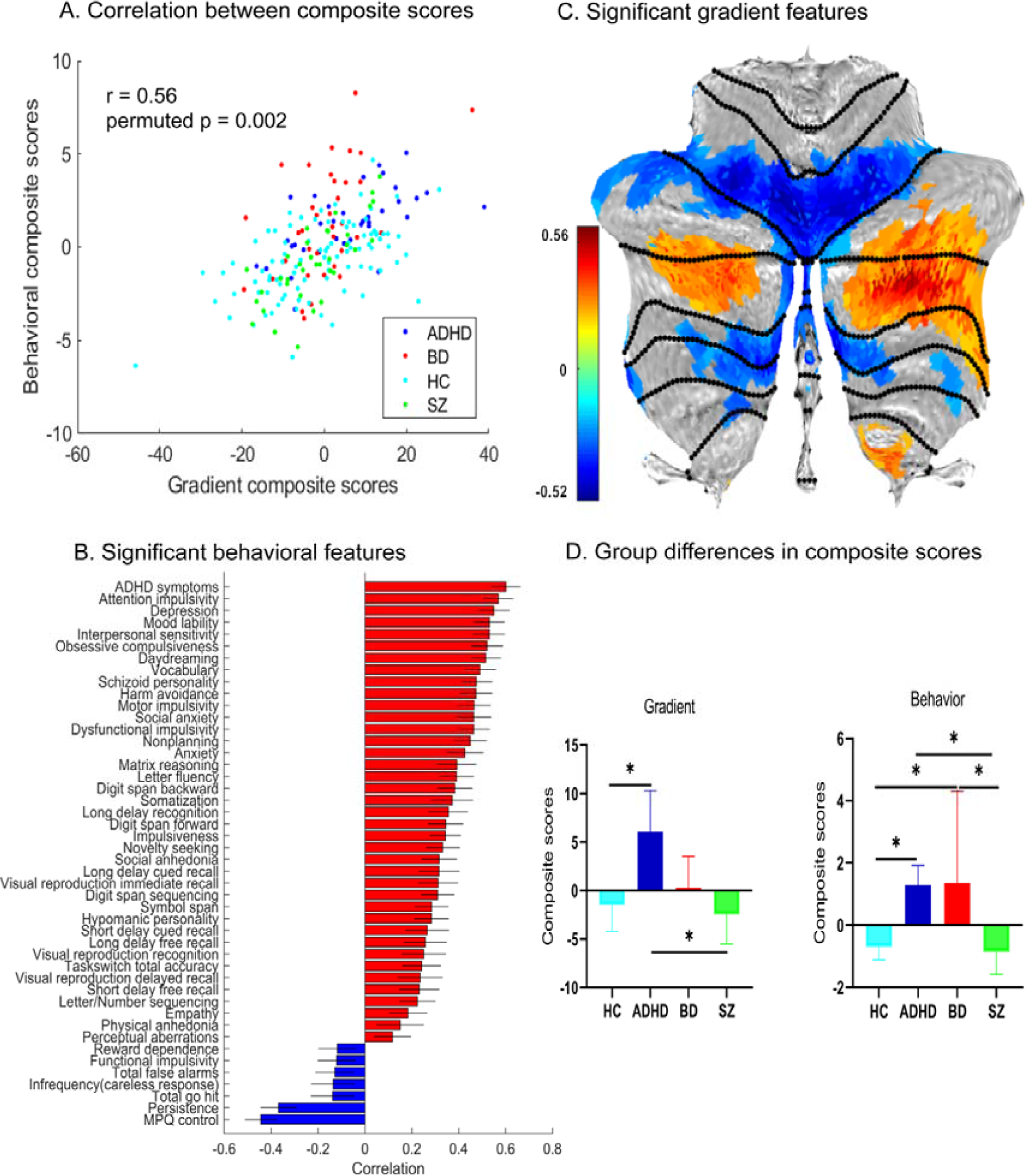
Latent variable 2: general lack of attention regulation. (A) Correlation between cerebellar connectivity gradient and behavioral composite scores of participants. (B) Significant behavioral features associated with LV2. The contribution of each behavior is measured by correlations between participants’ behavioral scores and the corresponding behavioral composite scores. Error bars indicate bootstrapped standard deviations. (C) Significant gradient pattern associated with LV2. The contribution of each voxel is measured by correlations between participants’ cerebellar gradient scores and the corresponding cerebellar gradient composite scores (FDR correction, q<0.05). (D) Group differences in cerebellar connectivity gradient and behavioral composite scores. Significant differences are indicated by asterisks (FDR correction, q < 0.05).

Furthermore, the four LVs were robust to GSR and total cerebellar grey matter volume regression, as indicated by the high correlation (r>0.83) between saliences of original PLS and PLS with GSR or total cerebellar grey matter volume regression. In addition, each diagnostic group contributed similarly to the overall composite correlations of these four LVs (FDR q > 0.05 for all pairwise comparisons, see Table S4). We also found that age, sex, education, site, or FD were not associated with any LV (Table S5).

### LV1: general psychopathology

The main contributors of behavior to LV1 were overall associated with greater psychopathology, e.g., higher impulsiveness, mood lability, dysfunctional impulsivity, anxiety, depression, somatization, social/physical anhedonia (Figure 2B) and psychotic symptoms (Table S6) including mania, delusions and hallucinations; in addition to worse high-order cognitive control (e.g., working memory). LV1 included positive weight in cerebellar lobules V, VI, VIIIA and VIIIB and negative weight in Crus I and II (Figure 2C). Notably, both cerebellar gradient and behavioral composite scores were higher in all diagnostic groups when compared with HCs (Figure 2D; all differences were statistically significant except for ADHD). Exploratory analyses indicated that higher cerebellar gradient and behavioral composite scores in LV1 were associated with greater medication load. There was no significant association between LV1 composite scores and substance use (Table S5). Our interpretation is that LV1 is associated mainly with general psychopathology and high-order cognitive control deficits (see discussion).

### LV2: general lack of attention regulation

The main contributors of behavior to LV2 were mainly involved in a general lack of attention regulation, e.g., higher ADHD symptoms, attention impulsivity, depression, mood lability, interpersonal sensitivity, daydreaming and social anxiety, and lower control ability and persistence (Figure 3B). LV2 included positive weight in cerebellar Crus I, II and lobule IX and negative weight in lobules VI, VIIB and VIIIA (Figure 3C). Notably, patients with ADHD had the highest cerebellar gradient scores for LV2 (Figure 3D). Behavioral composite scores were significantly higher in patients with ADHD or BD than in HC and patients with SZ. There was no significant association between composite scores and medication load or substance use (Table S5). Our interpretation is that LV2 is associated mainly with inadequate attention regulation (see discussion).

### LV3: internalizing symptoms

The main contributors of behavior to LV3 were mainly correlated with behavioral measures related to internalizing symptoms, e.g., higher harm avoidance, social anxiety, control, anhedonia, and somatization, and less externalizing symptoms, e.g., functional and motor impulsivity as well as novelty seeking (Figure 4B). LV3 included positive weight in cerebellar anterior vermis (I-VI) and negative weight in left Crus I, II, as well as lobules VIIIA and VIIIB (Figure 4C). Cerebellar gradient and behavioral composite scores were significantly higher in patients with BD or SZ when compared with patients with ADHD (Figure 4D). Higher cerebellar gradient and behavioral composite scores were associated with greater medication load (Table S5). There was no significant association between LV3 composite scores and substance use (Table S5). Our interpretation is that LV3 is associated mainly with higher internalizing symptoms and lower externalizing behavior (see discussion).

**Figure 4.**
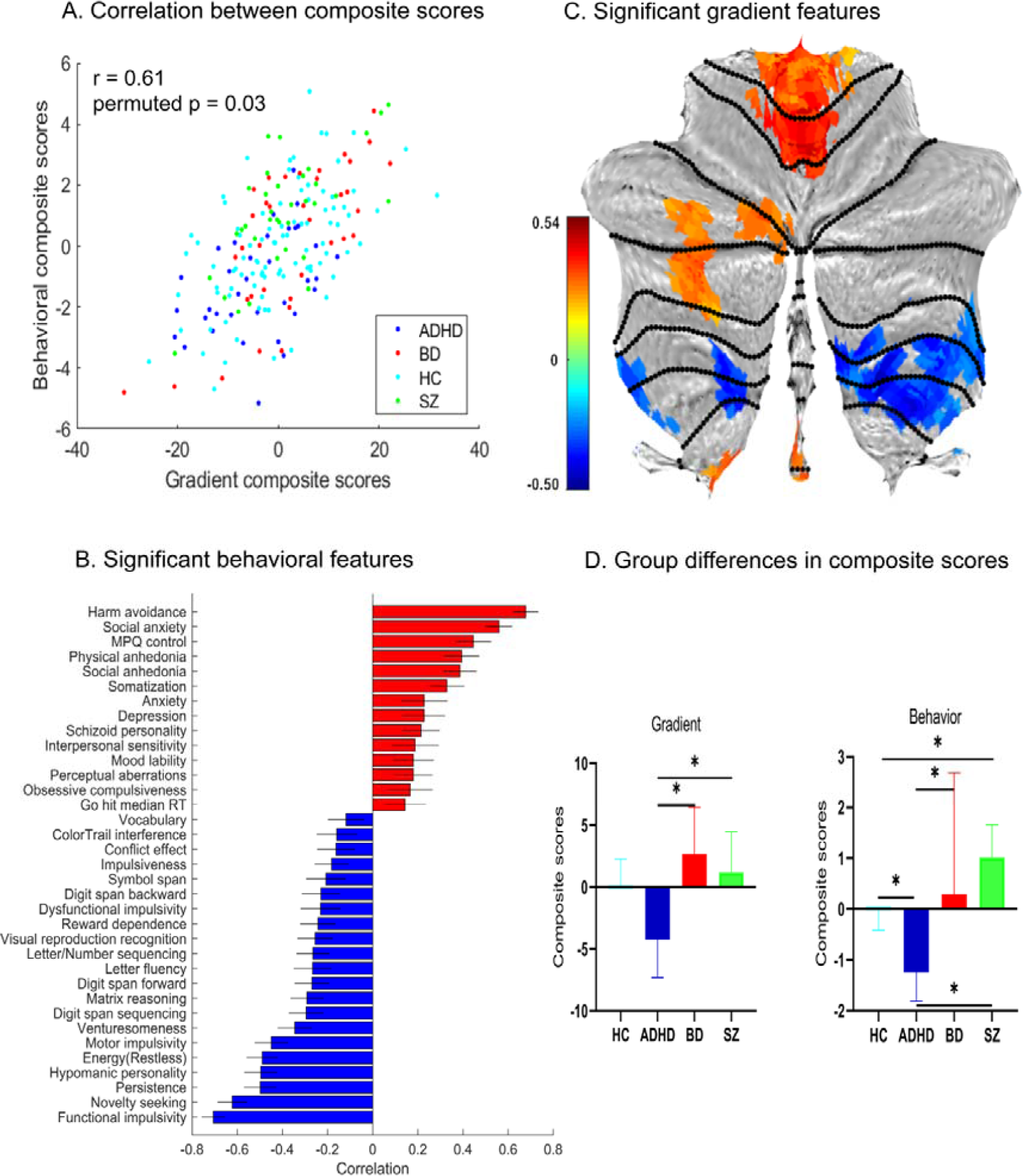
Latent variable 3: internalizing symptoms. (A) Correlation between cerebellar connectivity gradient and behavioral composite scores of participants. (B) Significant behavioral features associated with LV3. The contribution of each behavior is measured by correlations between participants’ behavioral scores and the corresponding behavioral composite scores. Error bars indicate bootstrapped standard deviations. (C) Significant gradient pattern associated with LV3. The contribution of each voxel is measured by correlations between participants’ cerebellar gradient scores and the corresponding cerebellar gradient composite scores (FDR correction, q<0.05). (D) Group differences in cerebellar connectivity gradient and behavioral composite scores. Significant differences are indicated by asterisks (FDR correction, q < 0.05).

### LV4: dysfunctional memory

The main contributors of behavior to LV4 included worse performance in multiple memory domains, as well as with less somatization, interpersonal sensitivity and depression (Figure 5B). LV4 included positive weight in Crus I, II and lobules IX and negative weight in lobule VI (Figure 5C). There was no significant difference among diagnostic groups (Figure 5D). There was no significant association between composite scores and medication load or substance use (Table S5). Our interpretation is that LV4 is associated mainly with dysfunctional memory (see discussion).

**Figure 5.**
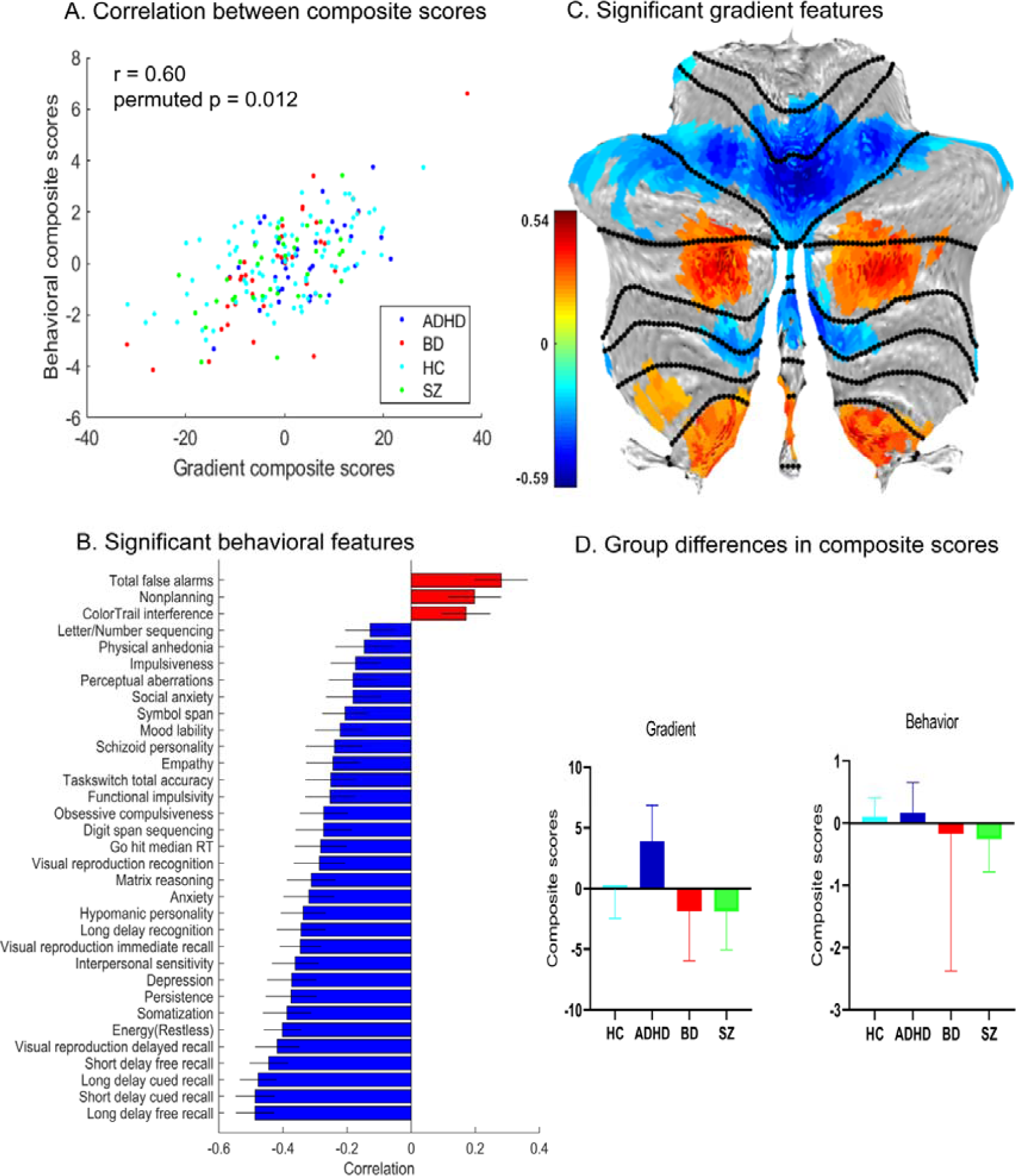
Latent variable 4: dysfunctional memory. (A) Correlation between cerebellar connectivity gradient and behavioral composite scores of participants. (B) Significant behavioral features associated with LV4. The contribution of each behavior is measured by correlations between participants’ behavioral scores and the corresponding behavioral composite scores. Error bars indicate bootstrapped standard deviations. (C) Significant gradient pattern associated with LV4. The contribution of each voxel is measured by correlations between participants’ cerebellar gradient scores and the corresponding cerebellar gradient composite scores (FDR correction, q<0.05). (D) Group differences in cerebellar connectivity gradient and behavioral composite scores. There were no significant differences among diagnostic groups in LV4 (FDR correction, q<0.05).

### Control Analyses

Additional control analyses ensured the robustness of the first four LVs to cerebellar gradients based on cerebellar-cerebral FC, confounding variables, non-Gaussian distributions of the behavioral data, diagnostic factors (HCs and patients separately), and site factors (each site separately) (see Supporting Information and Table S3). Results of PLS using only control individuals or only patients demonstrated moderate to high correlations with original saliences for the first four LVs. However, correlations dropped to 0.14 and 0.22 for LV5 (Table S3); hence we did not describe LV5.

## Discussion

Although the importance of cerebellar function in mental health and disease is increasingly recognized, the degree to which cerebellar connectivity is associated with transdiagnostic behavioral dimensions of psychopathology remains largely unknown. Leveraging a unique dataset including resting-state fMRI and behavioral assessments spanning clinical, cognitive, and personality traits, we found robust correlated patterns of cerebellar connectivity gradients and behavioral measures that could be represented in four transdiagnostic dimensions. Each dimension was associated with a unique spatial pattern of cerebellar connectivity gradients, and linked to different clusters of behavioral measures, supporting that individual variability in cerebellar functional connectivity can capture variability along multiple behavioral dimensions across psychiatric diagnoses. Our findings highlight the relevance of cerebellar neuroscience as a central piece for the study and classification of transdiagnostic dimensions of psychopathology – and ultimately for the diagnosis, prognosis, treatment, and prevention of mental illness. A large body of literature has shown cerebellar functional abnormalities in mental disorders.^2,3^

New trends in psychiatry focus on transdiagnostic dimensions of psychopathology.^4, 36^ The present study is the first to link both approaches. Adopting a transdiagnostic approach, three influential studies analyzing brain structure showed that alterations in cerebellar structure is associated with broad risk for psychopathology.^14–16^ However, these studies focused on clinical symptoms or cognitive function. The broader set of behavioral phenotypes in the present study allowed us to explore other dimensions of psychopathology, not constrained within the limits of clinical symptoms commonly investigated in many transdiagnostic studies.^15, 16, 28, 30, 37–39^ Prior cerebellar structure studies using factor analyses suggested the presence of latent dimensions of psychopathology such as internalizing symptoms, externalizing symptoms, and psychosis symptoms,^40^ as well as a general psychopathology (or p) factor.^41^ While these dimensions are reliable and reproducible, they are entirely derived from clinical assessments, not informed by brain-based data such as fMRI functional connectivity. More broadly, previous studies investigating functional connectivity-informed dimensions of psychopathology often ignore the importance of the cerebellum, e.g., by using a coarse delineation of the cerebellum with only a few regions of interest to represent the whole cerebellar information.^29, 30^ These limitations were overcome in the present investigation. Further, compared to methods that focus on a single view (such as factor analysis applied on clinical data), the present study derived behavioral dimensions from co-varying individual differences in connectivity gradients and behavioral measures. This approach resonates with the Research Domain Criteria research framework that encourages the integration of many levels of information.^36^

Our study indicates that individual variability in cerebellar functional connectivity gradient organization captures variability along multiple behavioral dimensions across mental health and disease. The associations with diverse dimensions of psychopathology were expected based on the consensus that the cerebellum is involved in virtually all aspects of behavior in health and disease.^1^ In 1998, Mesulam proposed that brain regions can be organized along a gradient ranging from sensory-motor to higher-order brain processes.^33^ In line with Mesulam, most of the variance of cerebellar RSFC resembles a gradient that spans from primary sensory-motor cortices to high–order transmodal regions of the default-mode network.^31^ This principal gradient may thus represent one fundamental principle driving a hierarchical organization of cerebellar motor, cognitive, and affective functions. Here we show for the first time that there is a link between this principal gradient of cerebellar organization and behavioral measures across individuals with and without diagnoses of cognitive or affective disease.

Functional gradient organizations in the brain have been proposed to reflect an architecture that optimizes the balance of externally and internally oriented functioning, which is critical for flexibility of human cognition.^33^ In this gradient organization, association areas are located at maximal distance from regions of primary areas that are functionally specialized for perceiving and acting in the here and now, supporting cognition and behavior not constrained by the immediate environment.^33, 42–44^ The intricate neuronal circuitry of the cerebellum has been hypothesized to function as a “forward controller,” creating internal models of how a given behavioral output will dynamically fit with contextual information,^45^ which is critical for monitoring and coordinating information processing in the service of mental processes.^1, 46, 47^ Thus, information processing in cerebellar circuits associated with multiple transdiagnostic dimensions of psychopathology shown here may reflect impaired monitoring and coordination of information—including one’s own thoughts and emotions—necessary to guide behavior, reflecting an imbalance of externally and internally oriented functioning, which may serve as possible intermediate phenotypes across mental health and diseases.

The most significant finding of the present investigation is the demonstration of an association between individual variations in cerebellar functional gradient values and multiple behavioral measures across mental health and diseases. Most behavior indicators were related to more than one dimension (Figure 2-5C). However, we noticed that the loadings of each behavior to each dimension can vary greatly, which highlighted the unique and different clusters of behavioral measures contributing to each dimension. As other brain-behavior association studies using multivariate analysis based on machine learnig,^48^ while it is not possible to provide a definitive characterization of the functional significance of each LV based on the analyses presented here, we here present one possible line of interpretation.

In LV1, greater behavioral composite score was associated with greater behavioral measures that we interpreted as general psychopathology and higher-cognitive control disabilities (including impulsiveness, mood lability, dysfunctional impulsivity, anxiety, depression, somatization, social/physical anhedonia and psychotic symptoms including mania, delusions and hallucinations). In line with the interpretation of LV1 as general psychopathology, both cerebellar gradient and behavioral composite scores were higher in all diagnostic groups when compared with HCs. Factor-analytic studies of multiple symptoms and diagnoses suggest that the structure of mental disorders can be summarized by three factors: internalizing, externalizing, and thought disorders.^40^ The empirical observation that even these three transdiagnostic latent factors are positively correlated^49^ has given rise to a more radical hypothesis, which is that there is the general psychopathology (or p) factor,^41^ which is thought to reflect individuals’ susceptibility to develop “any and all forms of common psychopathologies”.^50^ The p factor has been extended to index functional impairment, negative affect, emotion dysregulation, and cognitive deficits (e.g., attention and memory problems) (for a review see^4^). LV1 may thus reflect the p factor widely discussed in transdiagnostic cohorts.^41^

In LV2, greater behavioral composite scores were predominantly correlated with greater scores in areas related to a general lack of attention regulation including ADHD symptoms and attention impulsivity. Importantly, patients with ADHD had the highest gradient composite scores. LV2 might thus capture inattention and impulsivity/hyperactivity symptoms which characterize ADHD. However, other dimensions such as depression and schizoid personality were also included in LV2, arguing against a purely inattention-related nature of LV2.

In LV3, greater behavioral composite scores were dominantly correlated with greater behavioral measures related to internalizing symptoms (including harm avoidance, social anxiety, control, and anhedonia) and lower externalizing symptoms (including functional and motor impulsivity, novelty seeking, and hypomanic personality). LV3 may thus reflect an internalizing vs. externalizing factor.^40, 49^

LV4 was predominantly associated with negative correlations with behavioral measures, most strongly in the memory domain (long delay free recall, short delay cued recall, long delay cued recall, short delay free recall, and visual reproduction delayed recall). LV4 might thus dominantly reflect dysfunctional memory, although other behavioral domains also played a significant role in the behavioral composition of LV4 including restlessness, somatization, and persistence.

Notably, Kebets and colleagues investigated RSFC-informed dimensions of psychopathology in the CNP dataset,^29^ focusing on connectivity within and between cerebral and subcortical areas and derived a general psychopathology variable similar to LV1 in our study (other LVs were different), indicating that cerebral and cerebellar analyses might offer complementary information regarding the relationship between brain activity and behavioral measures. Future studies analyzing both cerebral and cerebellar data might determine whether cerebellar data offers similar or distinct information regarding the relationship between brain activity and behavioral measures when compared to analyses of cerebral data.

While providing novel evidence for associations between cerebellar hierarchical organization shown by fMRI and different dimensions of psychopathology, our analyses can provide only correlational – not causal – inferences between cerebellar function and behavior; future interventional experiments such as brain stimulation studies may be able to demonstrate not only an association but also a causal link between cerebellar function as indexed by functional gradients and behavioral measures. Another limitation that can be addressed in future research includes the relatively limited range of diagnostic categories in the patient population (ADHD, SZ, and BD); future research may extend our analyses to include additional patient populations. The analyses on the impact of medication and substance use were exploratory in our study; future studies with higher statistical power might adopt stronger statistical thresholds to study medication and substance use effects.

In summary, our results support an association between cerebellar functional connectivity gradients and multiple behavioral dimensions of psychopathology (general psychopathology, general lack of attention regulation, internalizing symptoms and dysfunctional memory) across healthy subjects and patients diagnosed with a variety of mental disorders. These findings highlight the importance of cerebellar function in transdiagnostic behavioral dimensions of psychopathology, and contribute to the development of cerebellar neuroscience as a tool that may significantly contribute to the diagnosis, prognosis, treatment, and prevention of cognitive and affective illness.

### Methods Participants

Data from the UCLA Consortium for Neuropsychiatric Phenomics (CNP) dataset ^34^ were downloaded from OpenNeuro (https://openneuro.org/datasets/ds000030/versions/00001). This dataset consists of neuroimaging and behavioral data from 272 right-handed participants, including both HC (n=130) and individuals with neuropsychiatric disorders including SZ (n=50), BD (n=49), and ADHD (n=43). Details about participant recruitment can be found in the original publication.^34^ Written informed consent was obtained from all participants and related procedures were approved by the Institutional Review Boards at UCLA and the Los Angeles County Department of Mental Health. Table 1 shows a summary of demographic and clinical information of the 198 participants who survived image preprocessing quality controls (see below).

### Behavioral assessment

The CNP behavioral measures encompass an extensive set of clinical, personality traits, neurocognitive and neuropsychological scores (Table S1). Behavioral measures were excluded from the partial least squares (PLS) analysis when data was missing for at least 1 participant among the 198 participants. As a result, we included a set of 55 behavioral and self-report measures from 19 clinical, personality traits, neurocognitive and neuropsychological tests in the PLS analysis. Table S2 summarized the behavioral measures for each group. Excluded behavioral measures were considered in post-hoc analyses (Table S6).

### Data Acquisition and Image Preprocessing

Resting-state functional and structural MRI data were collected on two 3T Siemens Trio scanners at UCLA using the same acquisition parameters. Resting-state functional MRI data were collected using a T2*-weighted echoplanar imaging sequence with the following scan parameters: TR/TE=2000ms/30 ms, flip angle = 90°, matrix 64 × 64, field of view (FOV) =192*192 mm^2^, 34 interleaved slices, slice thickness =4 mm, and oblique slice orientation.

The resting fMRI scan lasted 304 s for each participant, and 157 volumes were acquired. During scanning, all participants were instructed to keep relaxed and keep their eyes open. Additionally, T1-weighted high-resolution anatomical data were acquired for each participant using an MPRAGE sequence (scan parameters: TR/TE= 1900 ms/2.26 ms, matrix=256 × 256, FOV=250*250 mm^2^, sagittal plane, slice thickness =1 mm, 176 slices). The anatomical data were used to normalize functional data. See Supporting Information for details.

Among the 272 participants, there were seven participants with missing T1 weighted scans, four participants were missing resting-state functional MRI data scans, and 1 participant had signal dropout in the cerebellum,^51^ thus only data from 260 participants were preprocessed.

All preprocessing steps were consistent with our previous study.^52, 53^ In brief, the preprocessing steps included slice timing, realignment, normalization, wavelet despiking of head motion artifacts, regression of linear trend, Friston 24 head motion parameters, white matter and CSF signal, and filtering (0.01-0.1 Hz) (see Supporting Information for details). Because global signal may be an important neuroimaging feature in clinical populations,^54^ we did not conduct global signal regression (GSR) in our main analyses, but GSR was considered in control analysis. In addition, we excluded 42 participants due to head motion exceeding 1.5 mm or 1.5° rotation or with >10% images showing framewise displacements>0.5mm^55^ or mean FD>0.20mm during MRI acquisition. Further, we further excluded 20 participants because of incomplete coverage of the cerebellum. This process left 198 participants as a final sample for our study, among which there were 35 ADHD, 36 BD, 92 HC and 35 SZ participants.

### Cerebellar connectivity gradient extraction

We used diffusion map embedding ^56^ to identify a low-dimensional embedding gradient from a high-dimensional intra-cerebellar cortex connectivity matrix. Diffusion embedding results in multiple, continuous maps (“gradients”), which capture the similarity of each voxel’s functional connections along a continuous space. In other words, this data-driven analysis results in connectivity gradients that provide a description of the connectome where each voxel is located along a gradient according to its connectivity pattern. In order to maximize reliability, reproducibility, and interpretability, we only used the first gradient component in our analyses. The first gradient (or principal gradient) explains as much of the variance in the data as possible (∼30%, Figure 1), represents a well-understood motor-to-supramodal organizational principle in the cerebellar and cerebro-cerebral connections, and has been shown to be reproducible at the single subject level.^31^ (Guell et al., 2018; note that gradient 2 could not be reproduced as successfully as the principal gradient at the single-subject level) See Supporting Information for more details. We reported the intra-cerebellar FC gradient (6242 voxels) as the main result, but also included cerebellar-cerebral FC gradients in control analyses.

### Partial Least Squares analysis

We applied PLS to investigate the relationship between cerebellar connectivity gradient and behavioral measures across diagnostic categories. PLS is a multivariate statistical technique that derives latent variables (LVs), by finding weighted patterns of variables from two given data sets that maximally covary with each other.^57, 58^ Each LV is comprised of a cerebellar connectivity gradient pattern at voxel level (“gradient saliences”) and a behavioral profile (“behavioral saliences”). Individual-specific cerebellar gradient and behavioral composite scores for each LV were obtained by linearly projecting the gradient and behavioral measures of each participant onto their respective saliences. See Supporting Information for mathematical details. Because mean framewise displacement (FD) was negatively correlated with several behavioral measures and there were significant differences in age, sex, education, site, and mean FD across groups (Table 1), we regressed out these confounding effects from both behavioral and cerebellar gradient data before PLS analysis.

In order to evaluate the significance of the LVs, we applied permutation testing using 1000 permutations to determine the null distribution of the singular values. Considering significant group differences in various behavioral measures (Table S2), the permutation procedure was performed within each primary diagnostic group. Our results of interest were the top five LVs which explained at least 5% of covariance between cerebellar gradients and behavioral measures (see below). We applied a false discovery rate (FDR) correction of q < 0.05 on the permuted p-values of the five LVs to control for multiple comparisons.

To assess the contribution of a given gradient voxel or behavior to a given LV, we computed correlations between the original measure (gradient voxel or behavior) and the corresponding composite scores ^59, 60^. A large correlation (i.e., large weight, positive or negative) for a given measure (behavioral or gradient voxel) for a given LV indicates greater contribution of the behavior or gradient voxel to the LV. Then, the confidence intervals for these correlations were determined a by bootstrapping procedure that generated 500 samples with replacement from the original gradient and behavioral data. Considering significant diagnostic differences in many behavioral measures (Table S2), we took diagnostic groups into account within each bootstrap sample. To identify variables (gradient voxels or clinical measures) that make a significant contribution to the overall pattern, we calculated Bootstrapped Z scores as the ratio of each variable’s correlation coefficient (i.e., weight) to its bootstrap-estimated standard error. Then, we converted the Z scores to p values, which were FDR corrected (q<0.05).

To test the generalizability of each LV, we used a 10-fold cross-validation of the PLS analysis with 200 repetitions. Importantly, the cross-validation approach can help to guard against overfitting that arises from high dimensional neurobiological data.^35^ Specifically, first, we assigned 90% of the participants (within each primary diagnostic group) to the training set and the remaining 10% of participants (within each primary diagnostic group) to the test set. For each training set, PLS was used to estimate gradient and behavioral saliences (i.e., U_train_ and V_train_). Next, the test data were projected onto the gradient and behavioral patterns derived from the training set. This allowed us to estimate individual-specific gradient and behavior composite scores and their correlation for the test sample (i.e. corr(X_test_U_train_, Y_test_V_train_)) for LVs 1-4. This procedure was repeated 200 times to make sure the results are not biased by the initial split. Finally, we used a permutation test (behavioral data shuffled 1000 times within each diagnostic group) to assess the significance of these correlation coefficients.

Considering significant group differences in many behavioral measures (Table S2), we took diagnostic groups into account for the permutation procedure, bootstrapping procedure and cross-validation in the main text. However, when ignoring diagnostic groups (regarding all participants as one group), the results remained almost unchanged. See supplementary results for details.

If a given LV was statistically significant, we performed one-way ANOVA to test whether cerebellar gradient and behavioral composite scores of this LV were different among different diagnoses, if significant, least significant difference (LSD, in SPSS) post hoc tests were performed, which would help interpret the significant function of this LV. In addition, we furthermore tested whether the composite scores for significant LVs were correlated with confounding factors including age, sex, years of education, head motion, acquisition site, medication load (number of medications current use) and substance use(number of substances use, including nicotine, alcohol, cannabis and other psychotropic substances). T tests were performed for categorical variables, and Pearson’s correlations were performed for continuous measures. Given the exploratory nature of medication and substance use effect analysis in our study, we only consider the number of medications or substance current use, it should keep caution when interpreting these results. For binary measures, we used T tests, and for continuous measures, we used Pearson’s correlations. False discovery rate (FDR) correction (q<0.05) was applied to all analyses.

### Control Analyses

We tested whether LVs were robust to global signal regression, total cerebellar grey matter volume regression, cerebellar gradients based on cerebellar-cerebral FC, adding confounding variables (age, sex, education, site, and head motion) into the behavioral data for the PLS analysis, non-Gaussian distributions of the behavioral data, diagnostic factors (HCs and patients separately), and site factors (each site separately). To assess the robustness of each LV, we computed Pearson’s correlations between cerebellar gradient (or behavioral) saliences 24 obtained in each control analysis and cerebellar gradient (or behavioral) saliences from the original PLS analysis. Finally, to confirm that each diagnostic group contributed the same amount to the overall composite correlations, we used the Fisher r-to-z transformation to compare the pairwise r-values.^61^ See Supporting Information for details.

### Data and code availability

All data are freely provided by from the UCLA Consortium for Neuropsychiatric Phenomics (CNP) ^34^ available from OpenNeuro (https://openneuro.org/datasets/ds000030/versions/00001). Cerebellar connectivity gradients were constructed by BrainSpace toolbox (https://github.com/MICA-MNI/BrainSpace).^62^ We used the Matlab code from https://github.com/danizoeller/myPLS ^63^ and https://github.com/ThomasYeoLab/CBIG/tree/master/stable_projects/disorder_subtypes/Kebe ts2019_TransdiagnosticComponents,^29^ based on Krishnan et al. (2011)^58^ to implement the PLS calculating.

## Acknowledgements

This work was supported by the grant from National Key R&D Program of China (2018YFA0701400, C Luo), The grants from the National Nature Science Foundation of China (grant number: 61933003, D Yao, 81771822, C Luo and 81471634, C Luo), The Project of Science and Technology Department of Sichuan Province (2019YJ0179, C Luo), and the CAMS Innovation Fund for Medical Sciences (CIFMS) (No.2019-I2M-5-039, D Yao). We thank Dr Valeria Kebets, National University of Singapore for helpful comments and methodological discussion. We also thank the CNP investigators for making their data available for public access. All authors have agreed to this submission. A preprint of the present manuscript has been archived on the biorxiv.org preprint server (https://doi.org/10.1101/2020.06.15.153254).

## Supporting Information

### Supplemental Methods

#### Data acquisition and image preprocessing

MRI data were acquired two 3T Siemens Trio scanners, located at the Ahmanson-Lovelace Brain Mapping Center (Siemens version syngo MR B15) and the Staglin Center for Cognitive Neuroscience (Siemens version syngo MR B17) at UCLA. Resting-state functional MRI data were collected using a T2*-weighted echoplanar imaging (EPI) sequence with the following parameters: TR/TE=2000ms/30 ms, flip angle = 90°, matrix 64 × 64, field of view =192*192 mm^2^, 34 interleaved slices, slice thickness =4 mm, and oblique slice orientation. The resting fMRI scan lasted 304 s for each participant, and 157 volumes were acquired. Participants were asked to remain relaxed and keep their eyes open; they were not presented any stimuli or asked to respond during the scan. Additionally, T1-weighted high-resolution anatomical data were acquired for each participant using an MPRAGE sequence (scan parameters: TR/TE= 1900 ms/2.26 ms, matrix=256 × 256, FOV=250*250 mm^2^, sagittal plane, slice thickness =1 mm, 176 slices). The anatomical data were used to normalize functional data.

Among the 272 participants, there were seven participants with missing T1 weighted scans, four participants were missing resting-state functional MRI data scans, and 1 participant had signal dropout in the cerebellum,[1] thus only data from 260 participants were preprocessed. All preprocessing steps were carried out using the Data Processing & Analysis for (Resting-State) Brain Imaging (DPABI v4.1[2]) and Matlab scripts. Consistent with our previous study, [3,4] functional images were (1) discarded in the first five volumes, (2) slice-time corrected, (3) realigned, (4) co-registered to the high-resolution 3D anatomic volume, (5) warped into MNI152 standard space (resampling the voxel size into 3×3×3 mm^3^), (6) underwent wavelet despiking of head motion artifacts[5]), (7) underwent regression of motion and non-relevant signals, including linear trend, Friston 24 head motion parameters,[6,7] white matter (CompCor, 5 principal components), and CSF signal (CompCor, 5 principal components[8]), and (8) were filtered using a band-pass filter (0.01-0.1 Hz). Because global signal may be an important neuroimaging feature in clinical populations,[9] and global signal regression has been shown to induce anticorrelations in resting-state data,[10] we did not conduct global signal regression in our main analyses. Because the topic of global signal regression (GSR) is still controversial, we considered GSR in a separate control analysis. In addition, we excluded 48 participants due to head motion exceeding 1.5 mm or 1.5° rotation or with >10% frame-to-frame motion quantified by framewise displacements (FD>0.5mm, [11])) or mean FD > 0.20 mm during MRI acquisition. Further, we excluded 20 participants because of not full coverage of cerebellum. This process left 198 participants as a final sample of our study.

### Connectivity gradient analyses

In detail, the voxel-level connectivity matrix within cerebellar cortex for each subject was computed using Fisher Z-transformed Pearson correlations. Based on previous studies,[12–15] we thresholded the rsFC matrix with the top 10% of connections per row retained, whereas all others were zeroed. The negative connections were zeroed as well. Then, we used cosine distance to generate a similarity matrix that reflected the similarity of connectivity profiles between each pair of voxels.

Then, we used diffusion map embedding[16] to identify a low-dimensional embedding from a high-dimensional connectivity matrix. This methodological strategy has been able to successfully identify relevant aspects of functional organization in cerebral cortex and cerebellum in previous studies.[12,14] Similar to Principal Component Analysis (PCA), diffusion map embedding can identify principal gradient components accounting for the variance in descending order. If we applied PCA to the connectivity matrix, each voxel in cerebellar cortex would be assigned to a particular network with discrete borders. In contrast, diffusion map embedding allowed us to identify gradients of connectivity patterns from the similarity matrix. In this way, the result of diffusion embedding is not one single mosaic of discrete networks, but multiple, continuous maps (gradients), which capture the similarity of each voxel’s functional connections along a continuous space. In other words, this data-driven analysis results in connectivity gradients that provide a description of the connectome where each voxel is located along a gradient according to its connectivity pattern. Voxels with similar connectivity patterns are located close to one another along a given connectivity gradient. All gradients are orthogonal to each other and capture a portion of data variability in descending order.

There is a single parameter αto control how the density of sampling points affects the underlying manifold (α = 0, the maximal influence of sampling density; α = 1, no influence of sampling density) in the diffusion map embedding algorithm. Following previous studies,[12–14] we set α = 0.5, which can help retain the global relations between data points in the embedded space. Then, we used an average connectivity matrix calculated from all participants to produce a group-level gradient component template. We then performed Procrustes rotation to align the gradients of each participant to this template, following the strategy of previous analyses.[17] In order to maximize reliability, reproducibility, and interpretability, we only used the first gradient component in our analyses. The first gradient (or principal gradient) explains as much of the variance in the data as possible (∼30%, Figure 1), represents a well-understood motor-to-supramodal organizational principle in the cerebellar and cerebro-cerebral connections[14] and has been shown to be reproducible at the single subject level in the cerebellum (Guell et al., 2018; note that gradient 2 could not be reproduced as successfully as the principal gradient at the single-subject level). The explained variance of principle gradient (30%) was similar to recent reports using diffusion map embedding analyses in functional connectivity.[12–14,18]

We reported the intra-cerebellar FC gradient (6242 voxels) as the main results, but also we included the cerebellar-cerebral FC gradients in control analyses. The same calculation procedures used in intra-cerebellar functional connectivity gradient analysis were performed for cerebellar-cerebral cortical gradient analysis (cerebellar-cerebral cortical FC matrix).

### Partial least squares analysis

PLS is a multivariate procedure that seeks maximal correlations between two matrices by deriving LVs, which are optimal linear combinations of the original matrices.[19,20] We applied PLS to the cerebellar gradient and behavioral measures across diagnostic categories. Given two matrices, X_n*p_ and Y_n×q_, where n is the number of observations (e.g., participants, here n=198), p and q are the number of variables (e.g., cerebellar gradient (p=6242) and behavioral features (q=55), respectively), after z-scoring X and Y (across participants), we calculated the covariance matrix R=Y^T^X. Then, singular value decomposition (SVD) of R=USV^T^ produced in three low-dimensional latent variables: U and V are the singular vectors (called behavioral and cerebellar gradient saliences, similar to loadings in principal components analysis), while S is a diagonal matrix containing the singular values. After that, by linearly projecting the cerebellar gradient and behavioral measures of each participant onto their respective saliences, we obtained individual-specific cerebellar gradient and behavioral composite scores for each LV, which reflect the participants’ individual cerebellar gradient and behavioral contribution to each LV (similar to factor scores in principal components analysis). PLS seeks to find saliences that maximize the covariance between cerebellar gradient and behavioral composite scores. The covariance explained by each LV is estimated by dividing the squared singular value by the sum of all squared singular values. Because FD was negatively correlated with several behavioral measures mainly involving memory function (false discovery rate (FDR), q<0.05, including long delay free recall, visual reproduction immediate recall, delayed recall and recognition, matrix reasoning, and letter fluency) and there were significant differences in age, sex, education, site, and head motion (mean FD) across groups (Table 1), we regressed them out from both the behavioral and cerebellar gradient data before PLS analysis.

### Control Analyses Global signal regression

Given global signal regression is still the controversial issue in the rsfMRI field, in control analysis, we conducted global signal regression in the rsfMRI preprocessing to check whether the GSR significantly affects the four LVs.

### Regressing out cerebellar grey matter volume

To test whether the total cerebellar grey matter volume significantly affect the robustness of the four LVs, we re-computed PLS after regressing out total cerebellar grey matter volume from gradient features. We used the SUIT to calculate the total cerebellar grey matter volume.[24] Briefly, SUIT isolates the cerebellum and brainstem, then segments images into grey matter maps and normalizes these maps to a cerebellar template, ensuring superior cerebellar alignment across subjects. Normalized cerebellar grey matter maps were modulated by the Jacobian of the transformation matrix to preserve absolute grey matter volume. We extracted the summed modulated grey matter value (i.e., a measure of regional volume) for 28 cerebellar lobules defined in the probabilistic SUIT Atlas, and regarded resulting value as total cerebellar grey matter volume.[25]

### Cerebellar gradient based on cerebellar-cerebral FC

Given the cerebellar functional gradients can be similarly constructed based on intra-cerebellar FC or cerebellar-cerebral FC in the literature, we also tested cerebellar gradient based on cerebellar-cerebral FC. Intra-cerebellar connectivity gradient analysis focuses on exploring the intrinsic organization of the cerebellum without involving its connectivity profiles with the cerebral hemispheres or other brain structures. The cerebellar-cerebral cortical gradients emphasize the communication between cerebellar and cerebral cortex [14].

### Including confounds

Instead of regressing age, sex, education, site, and head motion (mean FD) out from the data prior to PLS analysis, we added them to the behavioral data for the new PLS analysis.

### Quantile normalization

Because many behavioral measures included in the PLS analysis were non-Gaussian distribution, to exclude the potential effect on the robustness of LVs, we used quantile normalization to improve the Gaussian distributions of the behavioral data and re-computed PLS between the normalized behavioral and cerebellar gradient data.

### Patients and sites factor

Furthermore, to ensure that our results were not separately driven by the HCs or by patients, we recomputed PLS using only control individuals or only patients. Finally, to ensure that results were not mainly driven by a single site, we recomputed PLS using data of each site separately.

### Contribution of each diagnostic group to the overall composite correlations

To confirm each diagnostic group contributes the same amount to the overall composite correlations, we used the Fisher r-to-z transformation to compare the pairwise r-values, i.e., correlation value between behavioral and gradient composite scores within each diagnostic group.[26]

## Supplemental Results

When ignoring diagnostic groups, i.e., regarding all participants as one group, the results remained almost unchanged. Specifically, the first four LVs were still significant (LV1: r=0.62, permuted p=0.008; LV2: r=0.56, permuted p=0.005; LV3: r=0.61, permuted p=0.038; LV4: r=0.60, permuted p=0.01). The significant behavioral and cerebral connectivity gradient features associated with each LV remained almost unchanged, see figure S2-S5. The 10-fold cross-validation for the first four LVs was still successful as indicated by significant correlation between cerebellar gradient and behavioral composite scores in the test folds (LV1, r=0.12, p=0.0029; LV2, r=0.16, p=0.0027; LV3, r=0.11, p=0.0029; LV4, r=0.07, p=0.0032).

### Control Analyses Global signal regression

Results were similar to the original PLS. Correlations between the saliences of the new and the original PLS analysis for the first four LVs ranged from 0.87 to 0.96 (Table S3), suggesting high correlation.

### Regressing out total cerebellar grey matter volume

Results were similar to the original PLS. Correlations between the saliences of the new and the original PLS analysis for the first four LVs ranged from 0.97 to 1 (Table S3), suggesting high correlation.

### Cerebellar gradient based on cerebellar-cerebral FC

When using cerebellar gradient based on cerebellar-cerebral FC, results were similar to the original PLS using the cerebellar gradient based on intra-cerebellar FC. Correlations between the saliences of the new and the original PLS analysis for the first four LVs ranged from 0.77 to 0.99, suggesting high correlation (Table S3).

### Including confounds

Results were similar to the original PLS, with moderate to high correlations between the saliences of the new and the original PLS analysis ranging from 0.61 to 0.93 for LVs 1-4 (Table S3).

### Quantile normalization

Results were similar to the original PLS, with high correlations between the saliences of the new and the original PLS analysis ranging from 0.95 to 0.99 for LVs 1-4 (Table S3).

### Patients and sites factor

When using healthy participants separately in the new PLS analysis, correlations between the saliences of the new and the original PLS analysis ranged between 0.46-0.83 for the first four LVs, suggesting moderate to high correlation. However, correlations dropped to 0.14 and 0.22 for LV5; hence we did not describe LV5 further. When considering only patients, correlations between the saliences of the new and the original PLS analysis ranged between 0.55-0.93 for the first four LVs, suggesting moderate to high correlation (Table S3).

When using only participants from site 1 in the new PLS analysis, correlations between the saliences of the new and the original PLS analysis ranged between 0.66-0.96 for the first four LVs, suggesting high correlation. When considering only participants from site 2, correlations between the saliences of the new and the original PLS analysis ranged between 0.49-0.97 for the first four LVs, suggesting moderate to high correlation (Table S3).

### Contribution of each diagnostic group to the overall composite correlations

There was no significant difference between pairs of correlation coefficients (Table S4, FDR q > 0.05 for all pairwise comparisons), suggesting that each diagnostic group contributed similarly to the overall composite correlations of these four LVs.

---

**Figure S1.**
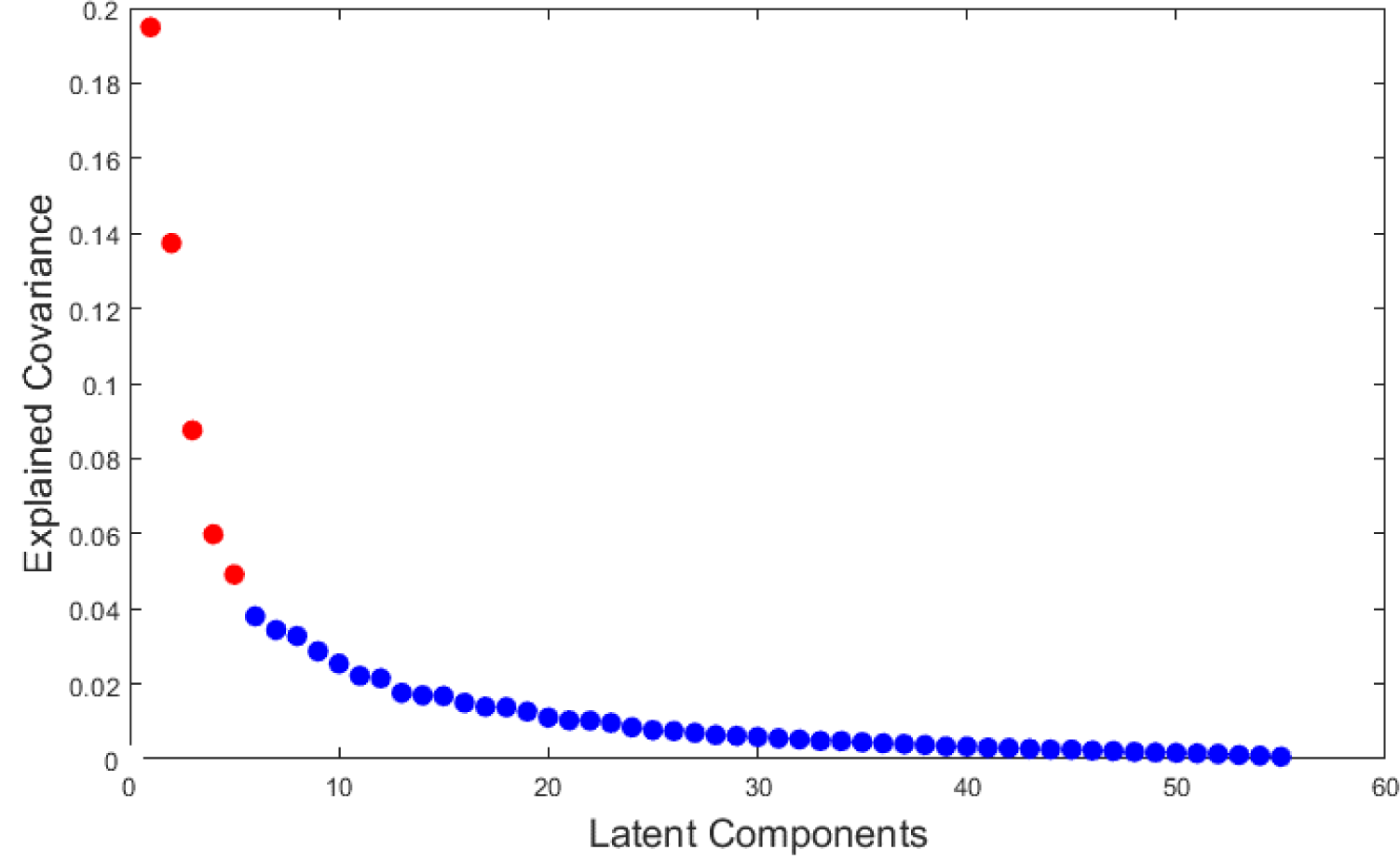
The amount of covariance explained by each LV. Five LVs survived after applying FDR correction (q<0.05) to the p-values derived from permutation tests.

**Figure S2.**
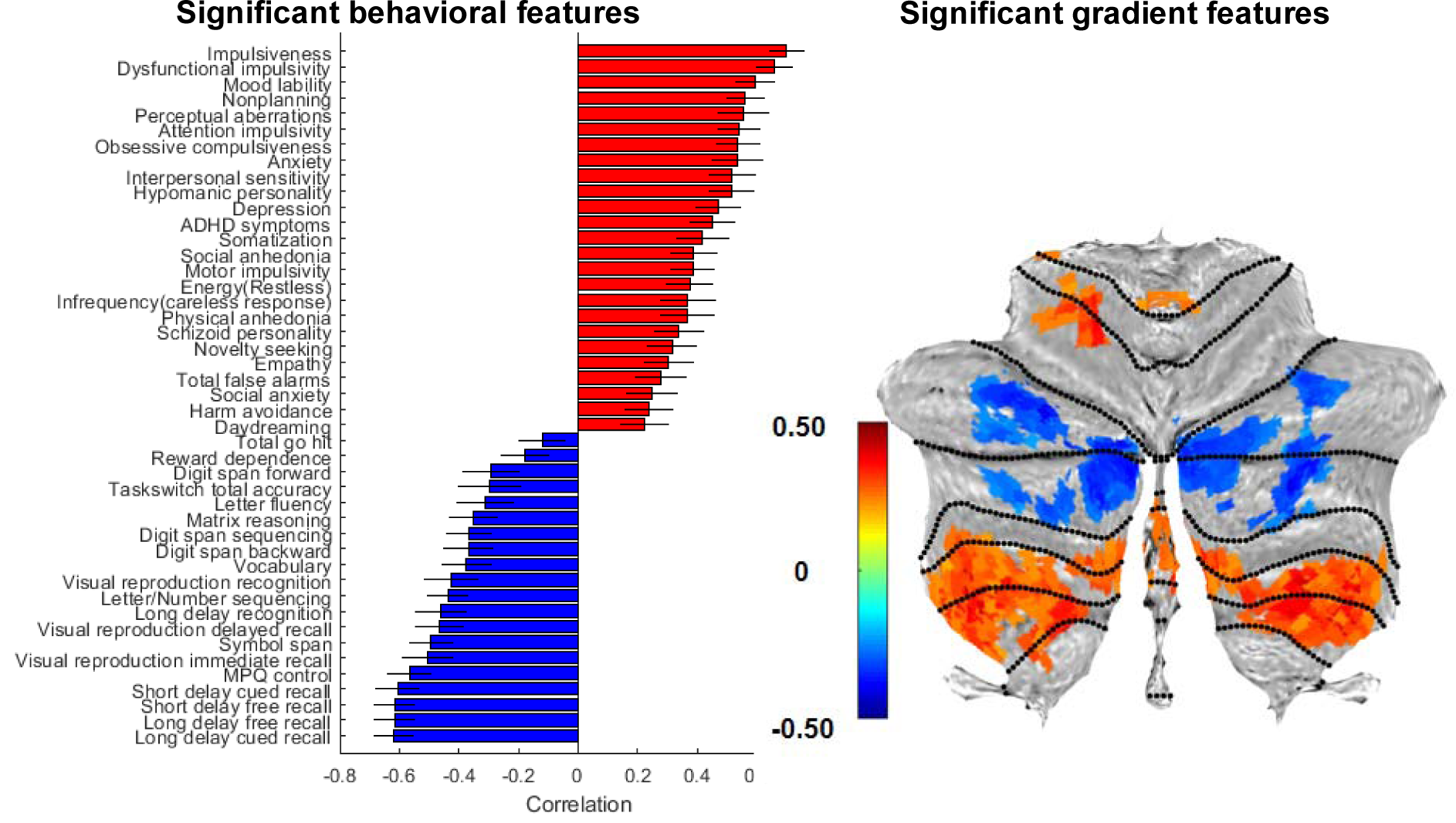
Significant behavioral and cerebral connectivity gradient features associated with LV1.

**Figure S3.**
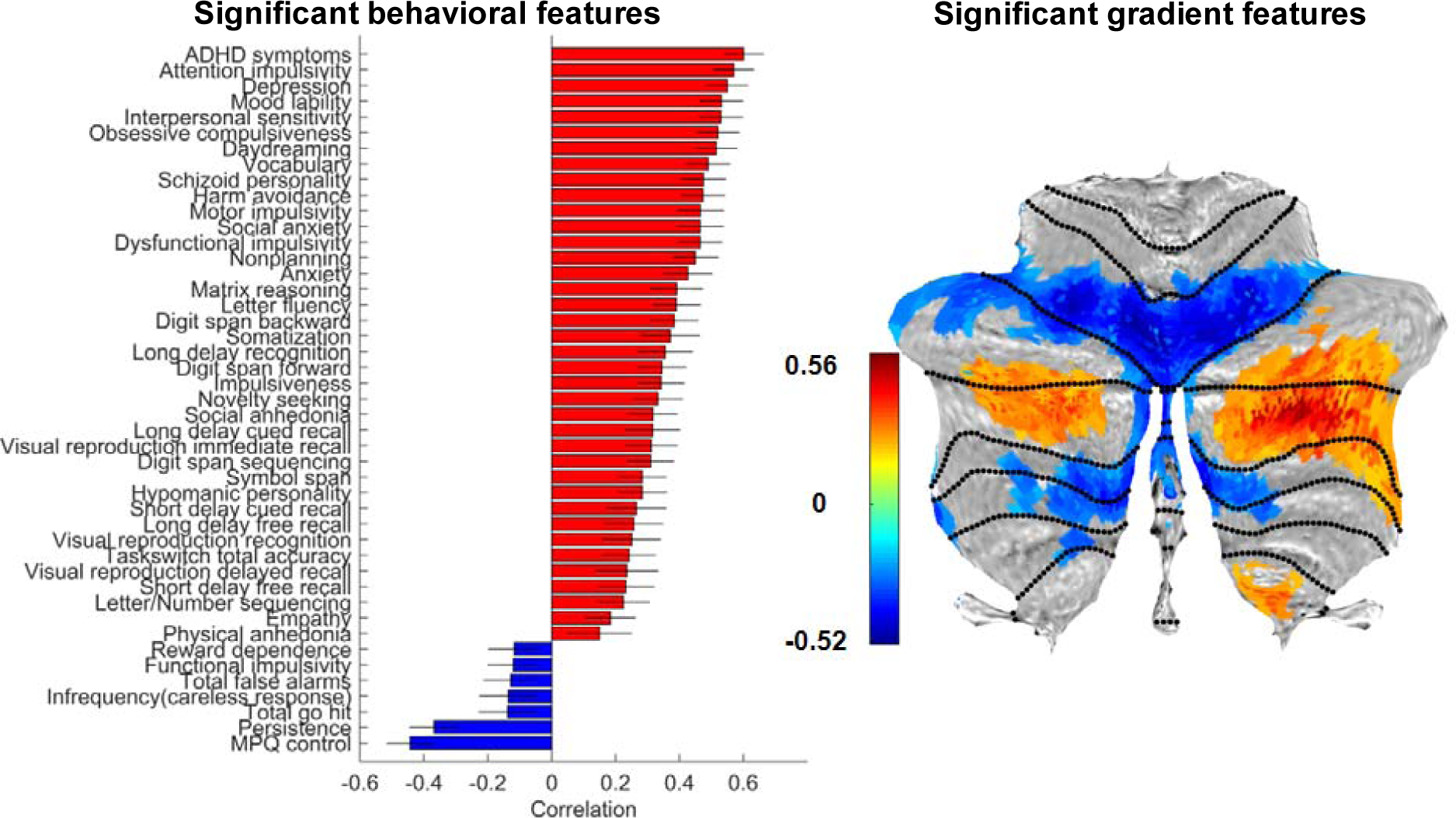
Significant behavioral and cerebral connectivity gradient features associated with LV2.

**Figure S4.**
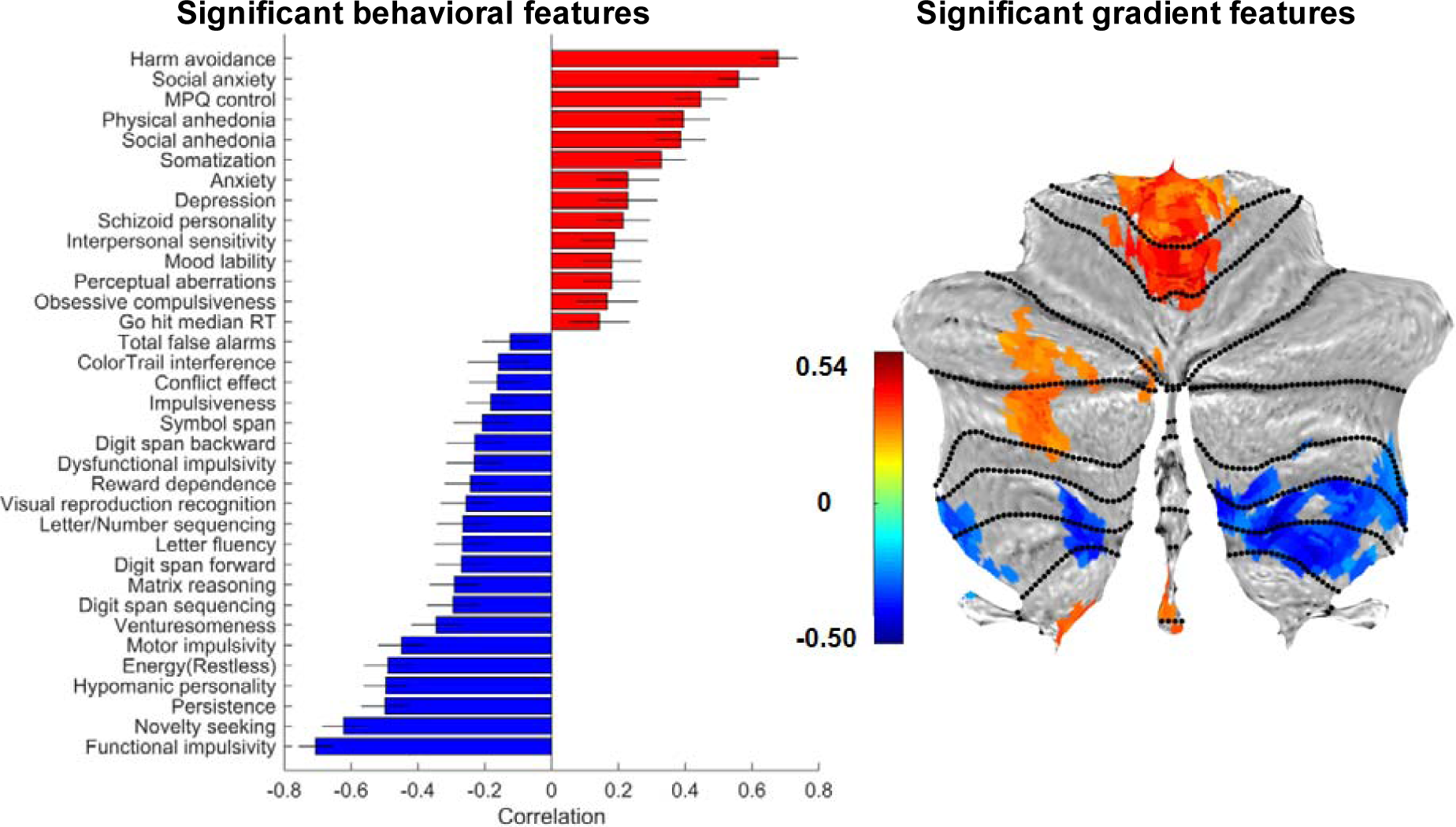
Significant behavioral and cerebral connectivity gradient features associated with LV3.

**Figure S5.**
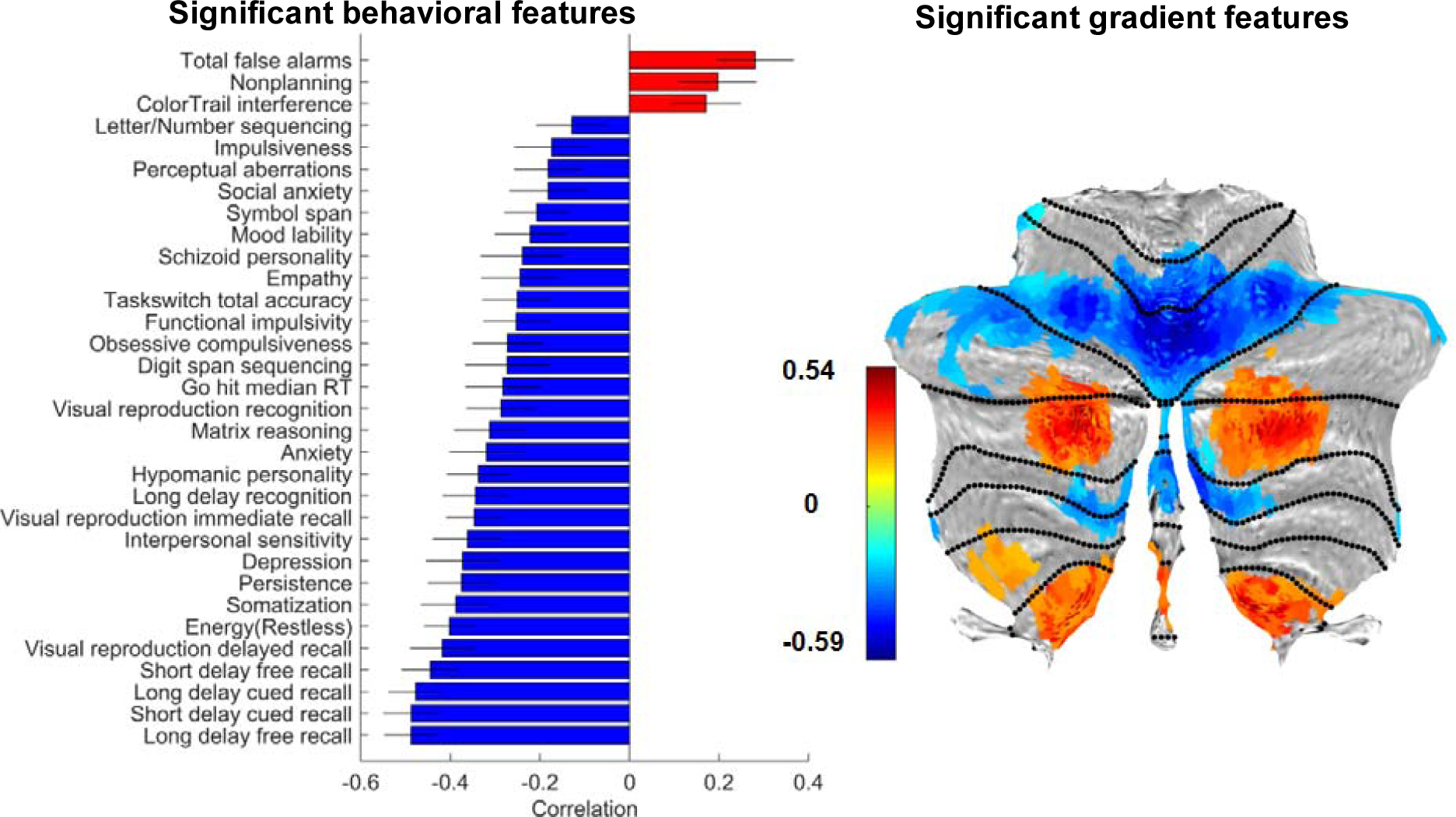
Significant behavioral and cerebral connectivity gradient features associated with LV4.

**Table S1.**
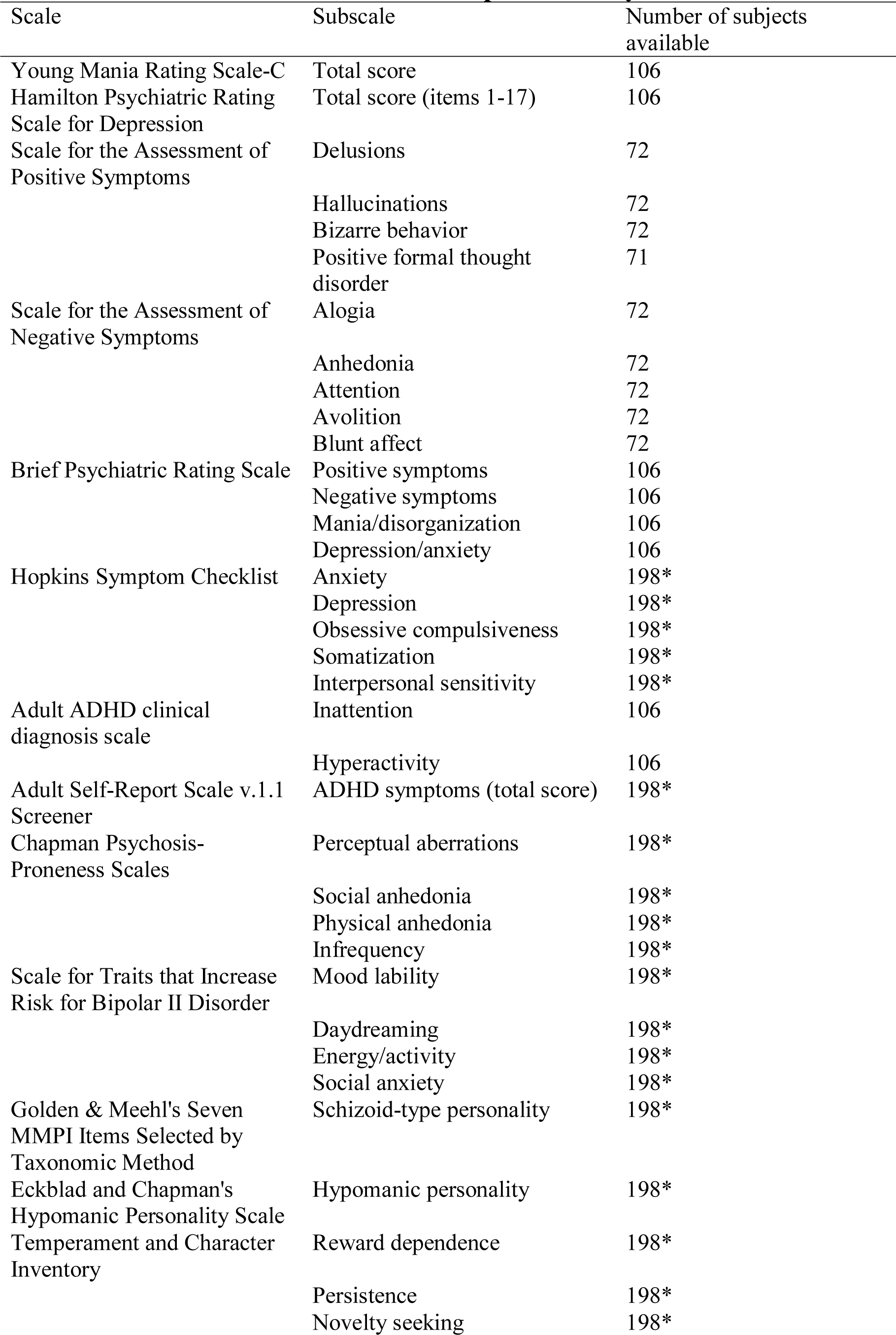
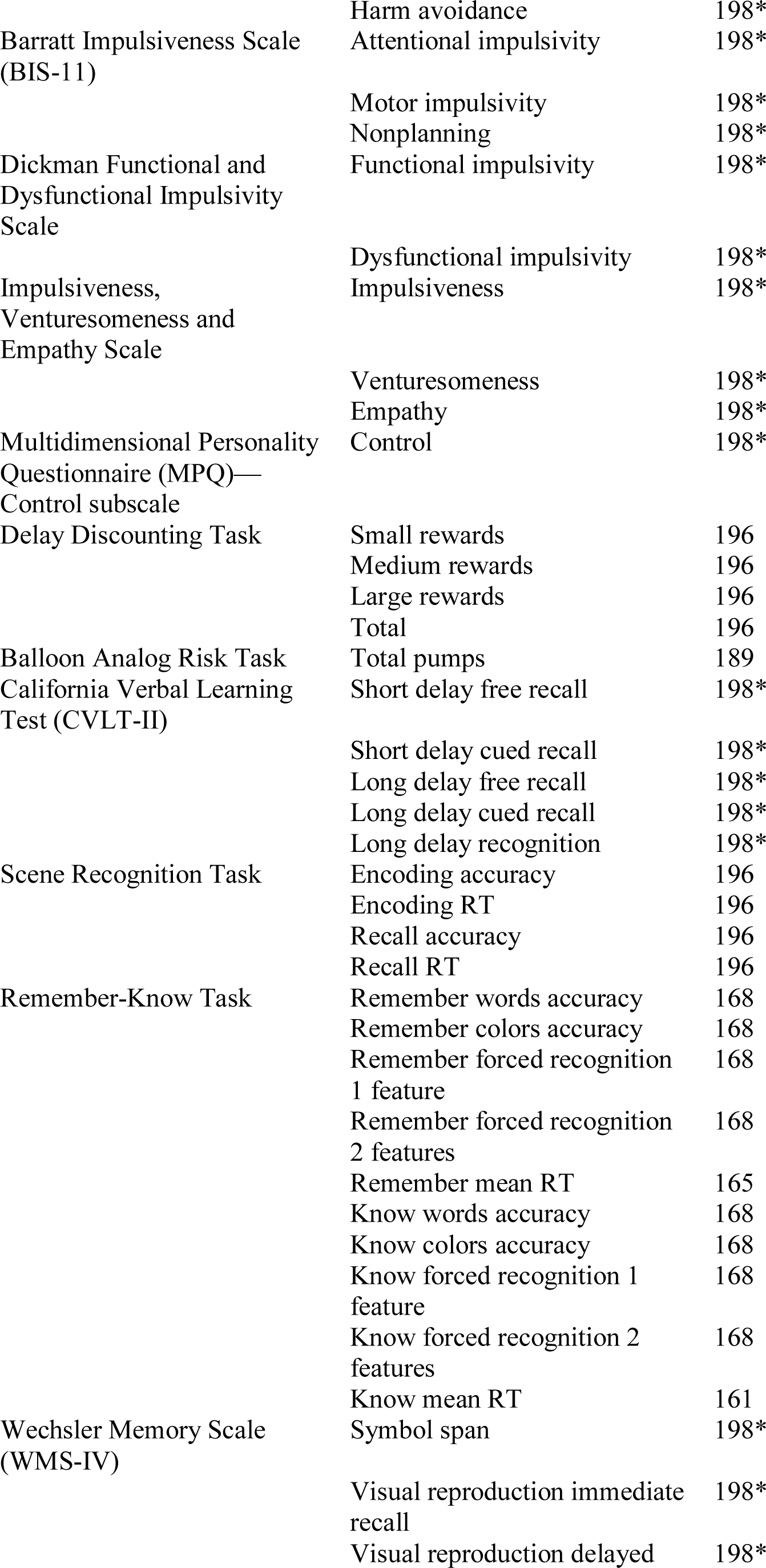
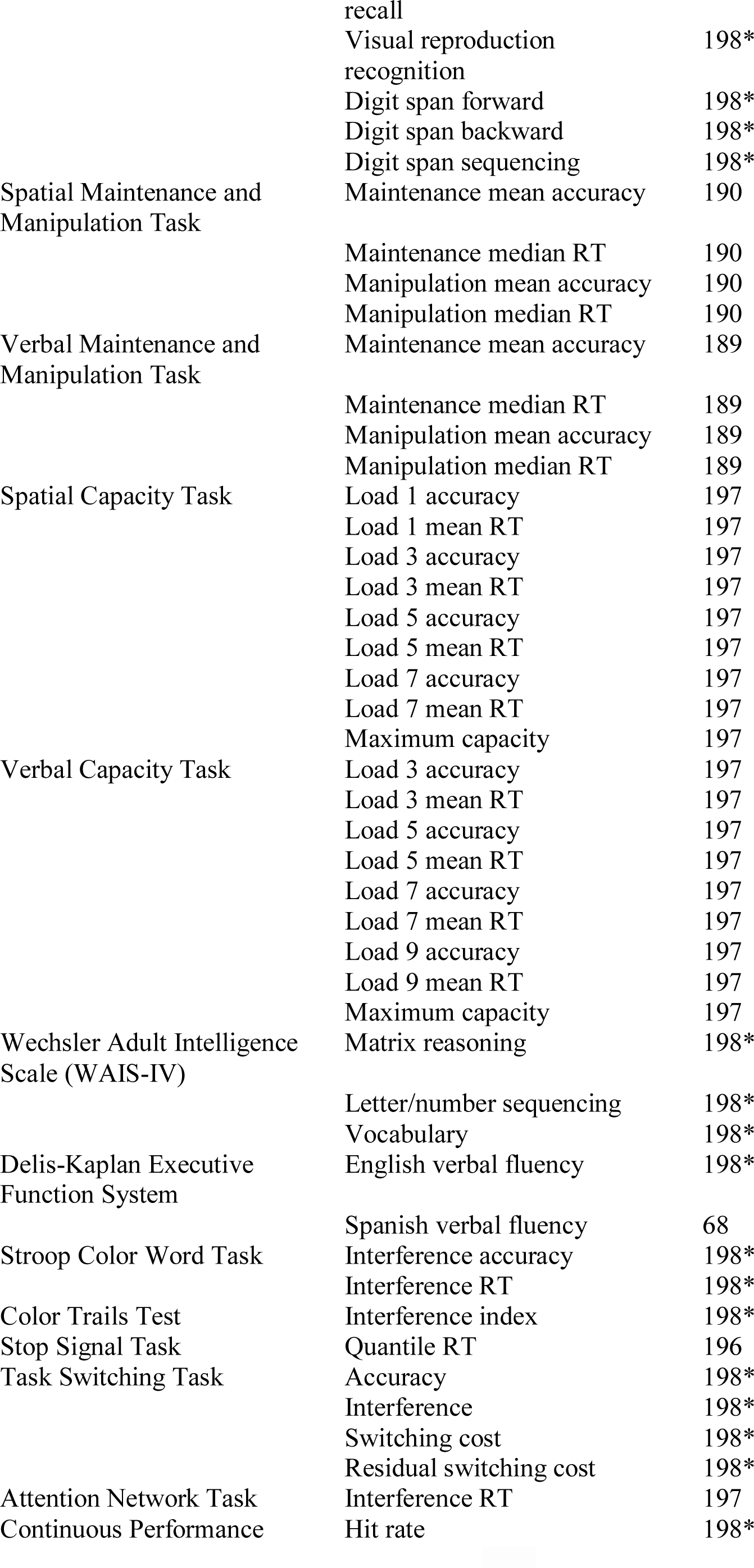
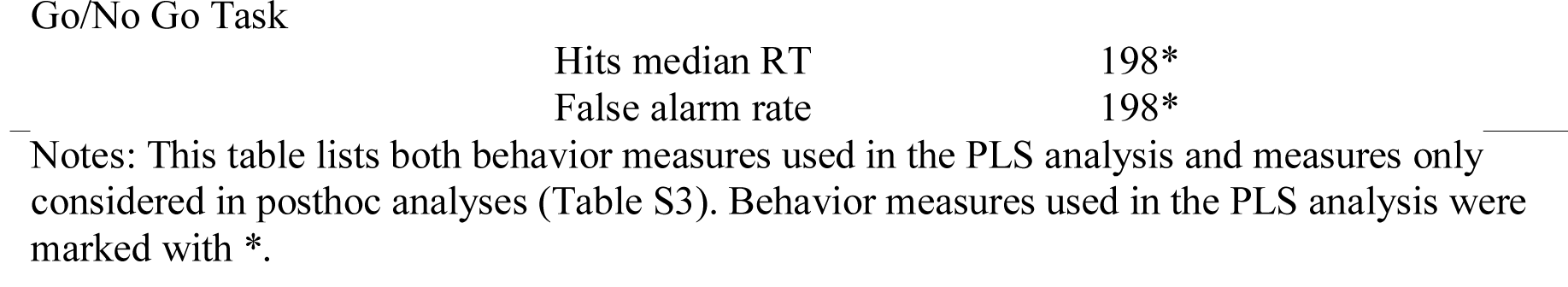
Behavior measures used in the present study

**Table S2.**
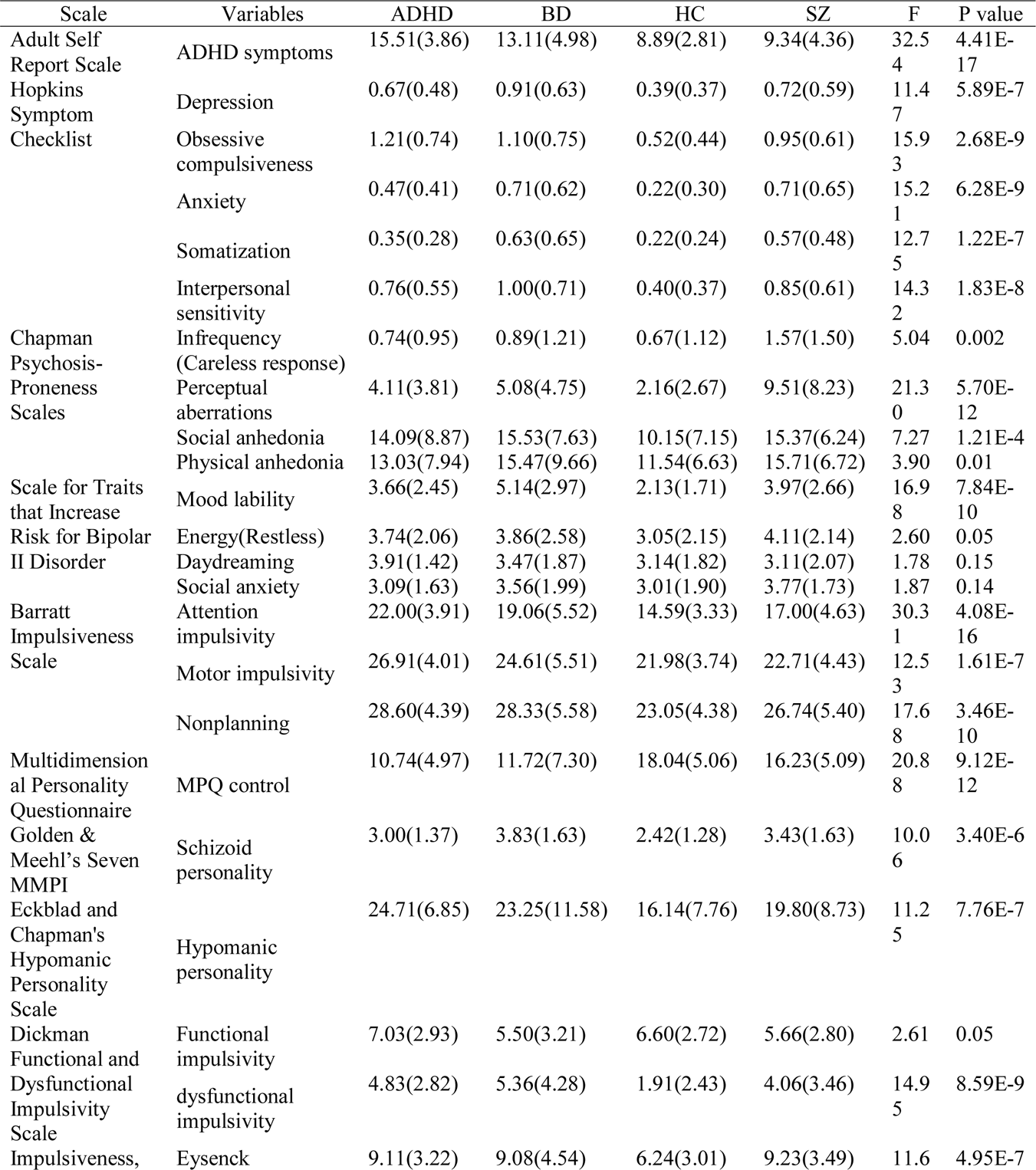
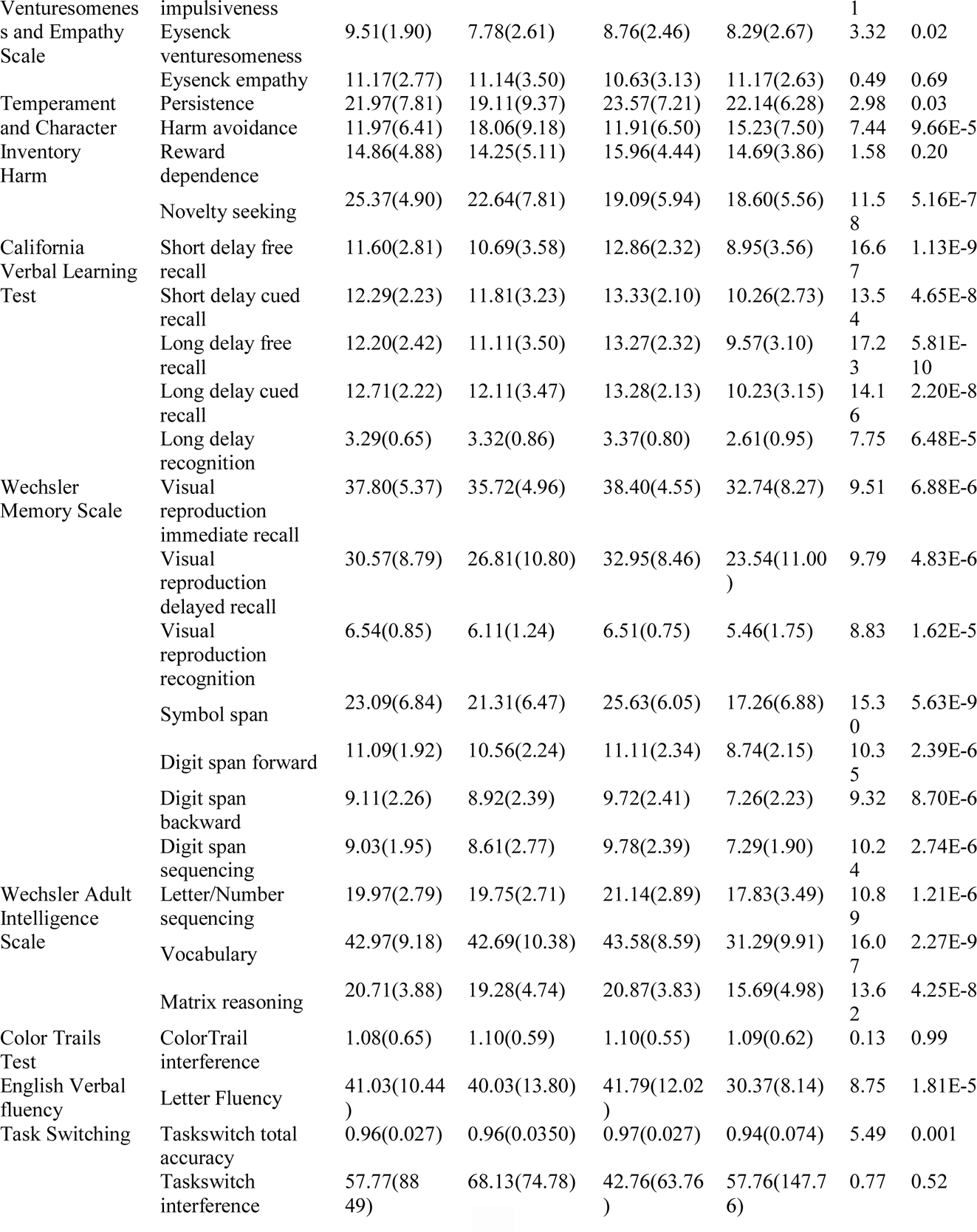
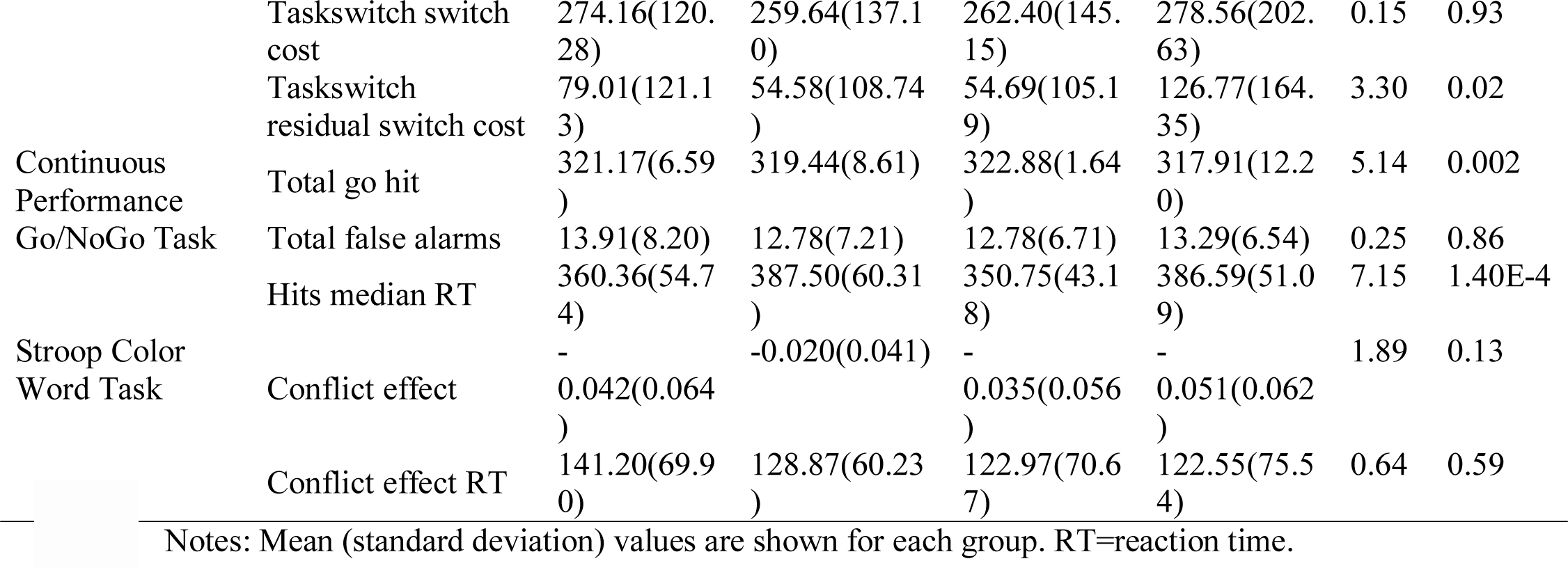
Group differences among the Fifty-five Behavioral measures in the PLS analysis

**Table S3.** Absolute correlations between cerebellar gradient (or behavioral) saliences obtained in control analyses and cerebellar gradient (or behavioral) saliences from the original PLS analysis.

|  | Latent<br>dimensi<br>on | GS<br>R | Regressing<br>out<br>cerebellar<br>grey matter<br>volume | Cerebella<br>r-<br>cerebral<br>Gradient | Confounds<br>included | Behavior<br>normalize<br>d | Contro<br>ls<br>Only | Patients<br>only | Site<br>#1 | Site<br>#2 |
| --- | --- | --- | --- | --- | --- | --- | --- | --- | --- | --- |
| Correlations<br>with<br>original<br>gradient<br>salience | LV #1 | 0.9<br>1 | 1 | 0.83 | 0.77 | 0.99 | 0.46 | 0.61 | 0.76 | 0.68 |
|  | LV #2 | 0.9<br>2 | 0.97 | 0.85 | 0.85 | 0.98 | 0.70 | 0.55 | 0.79 | 0.76 |
|  | LV #3 | 0.8<br>7 | 0.98 | 0.77 | 0.89 | 0.97 | 0.66 | 0.77 | 0.75 | 0.49 |
|  | LV #4 | 0.9<br>1 | 0.99 | 0.81 | 0.62 | 0.97 | 0.57 | 0.67 | 0.66 | 0.69 |
|  | LV #5 | 0.8<br>3 | 1 | 0.82 | 0.58 | 0.95 | **0.14** | 0.55 | 0.52 | 0.45 |
| Correlations<br>with<br>original<br>behavioral<br>salience | LV #1 | 0.9<br>8 | 1 | 0.99 | 0.93 | 1 | 0.83 | 0.82 | 0.96 | 0.97 |
|  | LV #2 | 0.9<br>5 | 0.98 | 0.96 | 0.84 | 0.97 | 0.80 | 0.59 | 0.90 | 0.89 |
|  | LV #3 | 0.9<br>6 | 0.99 | 0.93 | 0.93 | 0.97 | 0.87 | 0.93 | 0.89 | 0.70 |
|  | LV #4 | 0.9<br>4 | 0.99 | 0.92 | 0.61 | 0.95 | 0.71 | 0.64 | 0.74 | 0.76 |
|  | LV #5 | 0.8<br>7 | 1 | 0.94 | 0.63 | 0.95 | **0.22** | 0.69 | 0.63 | 0.61 |

**Table S4.** Comparisons between pairs of correlation coefficients between gradient composite scores and behavioral composite scores

| LV1(separated group-1) | LV2(separated group-1) | LV3(separated group-1) | LV4(separated group-1) |
| --- | --- | --- | --- |
| r_hc=0.6099;<br>r_patients=0.5696<br>z = 0.4272, p = 0.6692 | r_hc=0.5415;r_patients=0.5943<br>z = -0.5390, p = 0.5899 | r_hc=0.5548;r_patients =0.6571<br>z = -1.1222, p = 0.2618 | r_hc=0.5466;r_patients=0.6667<br>z = -1.3216, p = 0.1863 |
| LV1(separated group-2) | LV2(separated group-2) | LV3(separated group-2) | LV4(separated group-2) |
| r_hc=0.6099;<br>r_sz=0.4539<br>r_bd=0.7114;<br>r_adhd=0.5740<br>hc-sz: z = 1.0633, p = 0.2877;<br>hc-bd: z = -0.8893, p = 0.3738<br>hc-adhd: z = 0.2683, p = 0.7885<br>sz-bd: z = -1.6139, p = 0.1065<br>sz-adhd: z = -0.6555, p = 0.5122<br>bd-adhd: z = 0.9534, p = 0.3404 | r_hc=0.5415; r_sz=0.7511<br>r_bd=0.5103; r_adhd=0.6037<br>hc-sz: z = -1.7912, p = 0.0733<br>hc-bd: z = 0.2117, p = 0.8324<br>hc-adhd: z = -0.4496, p = 0.6530<br>sz-bd: z = 1.6620, p = 0.0965<br>sz-adhd: z = 1.1061, p = 0.2687<br>bd-adhd: z = -0.5474, p = 0.5841 | r_hc=0.5548;<br>r_sz=0.6782<br>r_bd=0.7275;<br>r_adhd=0.4116<br>hc-sz: z = -0.9727, p = 0.3307<br>hc-bd: z = -1.4627, p = 0.1436<br>hc-adhd: z = 0.9109, p = 0.3624<br>sz-bd: z = -0.3935, p = 0.6940<br>sz-adhd: z = 1.5529, p = 0.1204<br>bd-adhd: z = 1.9583, p = 0.0502 | r_hc=0.5466; r_sz=0.5607<br>r_bd=0.8036; r_adhd=0.4845<br>hc-sz: z = -0.0986, p = 0.9214<br>hc-bd: z = -2.4296, p = 0.0151<br>hc-adhd: z = 0.4108, p = 0.6812<br>sz-bd: z = -1.9139, p = 0.0556<br>sz-adhd: z = 0.4200, p = 0.6745<br>bd-adhd: z = 2.3372, p = 0.0194 |

Notes: There was no significant difference between pairs of correlation coefficients (FDR q > 0.05 for all pairwise comparisons)

**Table S5.** Associations between cerebellar gradient or behavior composite scores and confounding factors

|  | LV1 |  |  |  | LV2 |  |  |  | LV3 |  |  |  | LV4 |  |  |  |
| --- | --- | --- | --- | --- | --- | --- | --- | --- | --- | --- | --- | --- | --- | --- | --- | --- |
|  | Gradient composite scores |  | Behavioral composite scores |  | Gradient composite scores |  | Behavioral composite scores |  | Gradient composite scores |  | Behavioral composite scores |  | Gradient composite scores |  | Behavioral composite scores |  |
|  | r/t | p | r/t | p | r/t | p | r/t | p | r/t | p | r/t | p | r/t | p | r/t | p |
| Age | 0 | 1 | 0 | 1 | 0 | 1 | 0 | 1 | 0 | 1 | 0 | 1 | 0 | 1 | 0 | 1 |
| Sex | 0 | 1 | 0 | 1 | 0 | 1 | 0 | 1 | 0 | 1 | 0 | 1 | 0 | 1 | 0 | 1 |
| Education | 0 | 1 | 0 | 1 | 0 | 1 | 0 | 1 | 0 | 1 | 0 | 1 | 0 | 1 | 0 | 1 |
| Site | 0 | 1 | 0 | 1 | 0 | 1 | 0 | 1 | 0 | 1 | 0 | 1 | 0 | 1 | 0 | 1 |
| Motion | 0 | 1 | 0 | 1 | 0 | 1 | 0 | 1 | 0 | 1 | 0 | 1 | 0 | 1 | 0 | 1 |
| Total brain volume | -0.04 | 0.59 | -0.08 | 0.26 | -0.03 | 0.63 | 0.05 | 0.48 | -0.05 | 0.44 | -0.07 | 0.31 | 0.02 | 0.80 | 0.04 | 0.62 |
| Cerebellar volume | -0.10 | 0.20 | -0.04 | 0.55 | 0.15 | 0.03 | 0.17 | 0.02 | -0.06 | 0.38 | -0.10 | 0.18 | 0.07 | 0.32 | 0.02 | 0.73 |
| Medication load | **0.35** | **4.7E-7** | **0.39** | **1.2E-8** | -0.09 | 0.19 | 0.07 | 0.30 | **0.18** | **0.01** | **0.28** | **6.2E-5** | -0.07 | 0.30 | 0.06 | 0.43 |
| Substance use | 0.12 | 0.08 | 0.10 | 0.16 | 0.07 | 0.35 | 0.15 | 0.03 | -0.13 | 0.06 | -0.10 | 0.15 | -0.03 | 0.65 | -0.07 | 0.35 |

Notes: T tests were performed for binary measures, and Pearson's correlations were performed for continuous measures. Bold refers to significant associations that survived FDR correction ( $q < 0.05$ ).

**Table S6.**
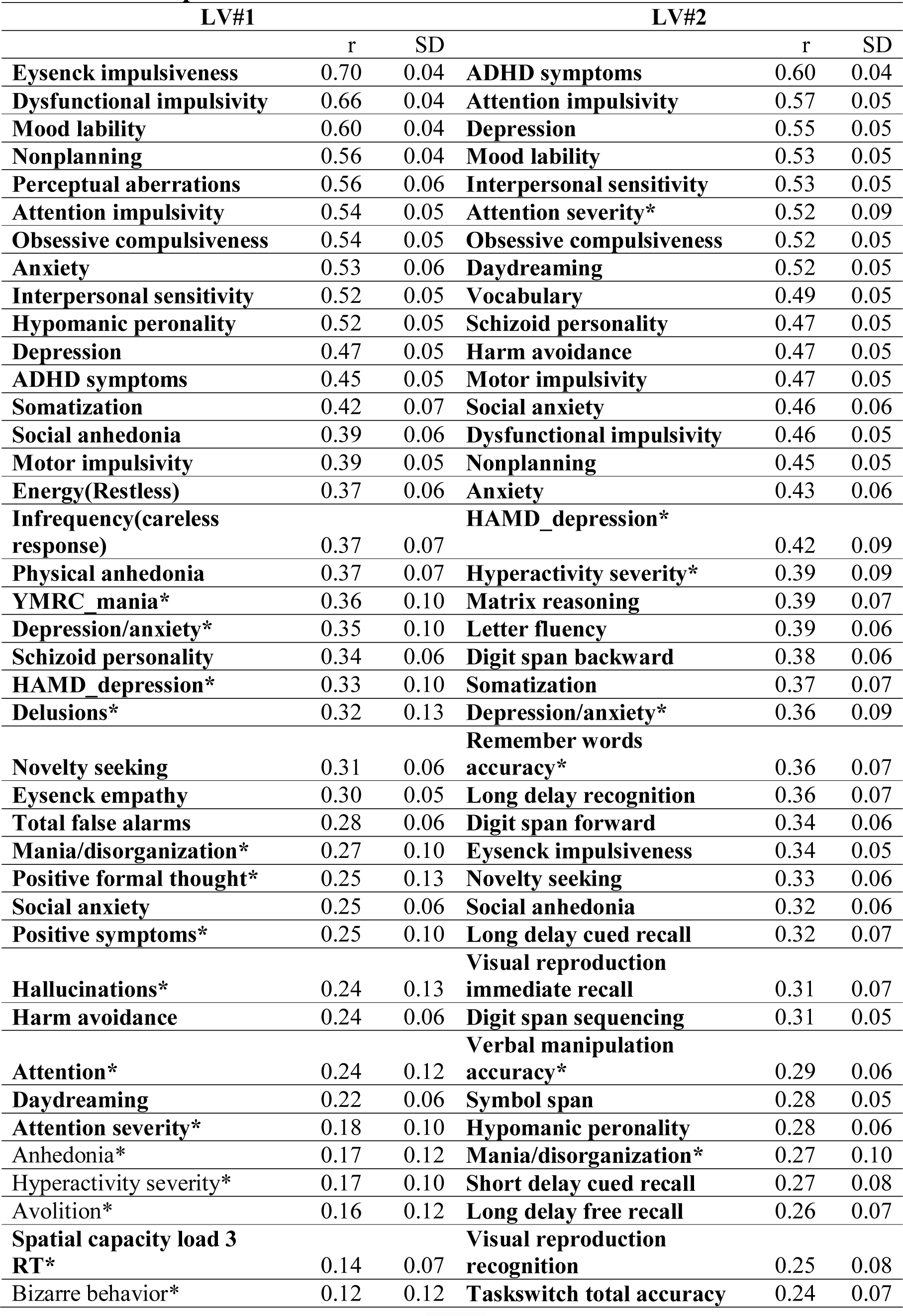
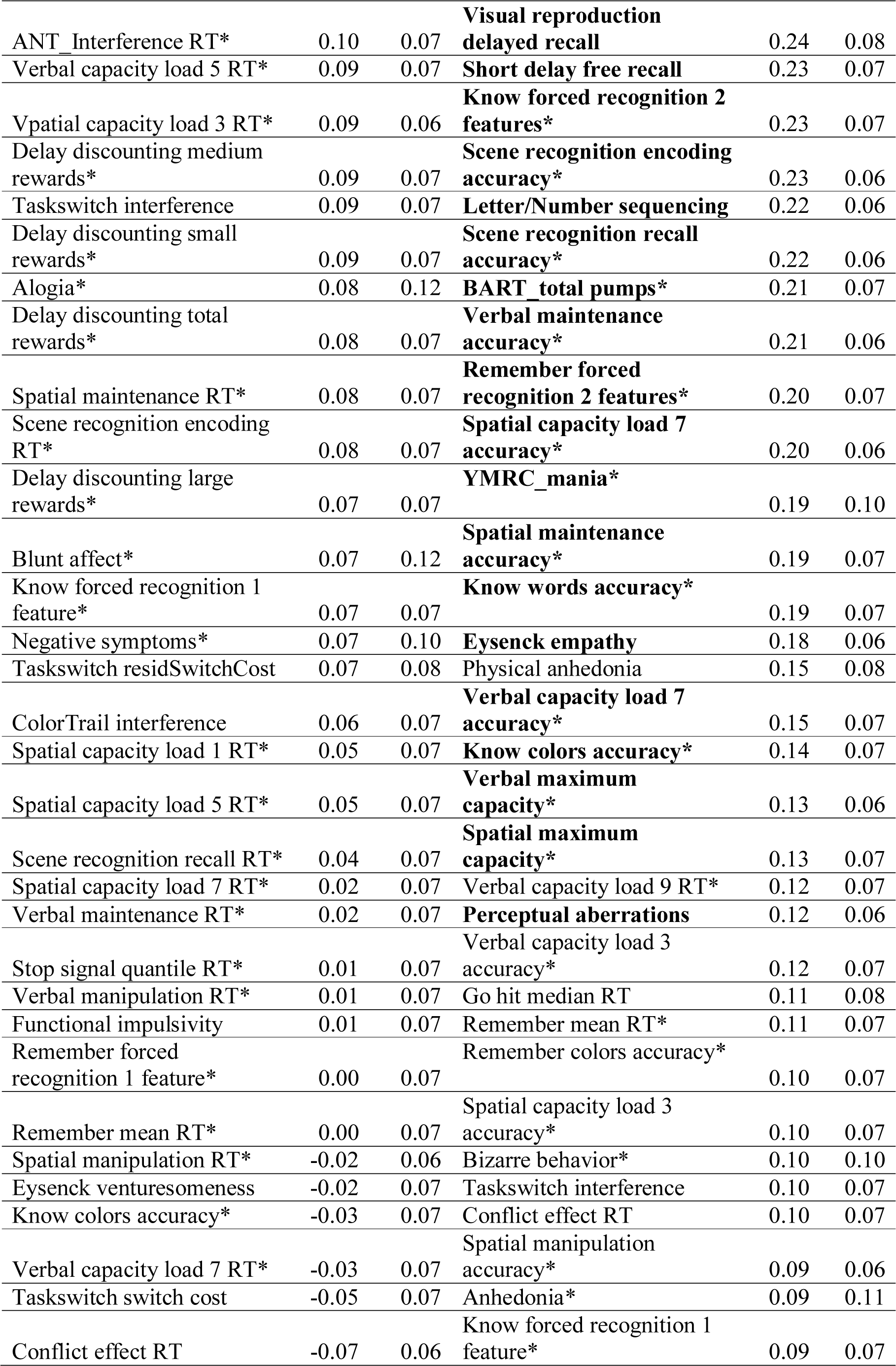
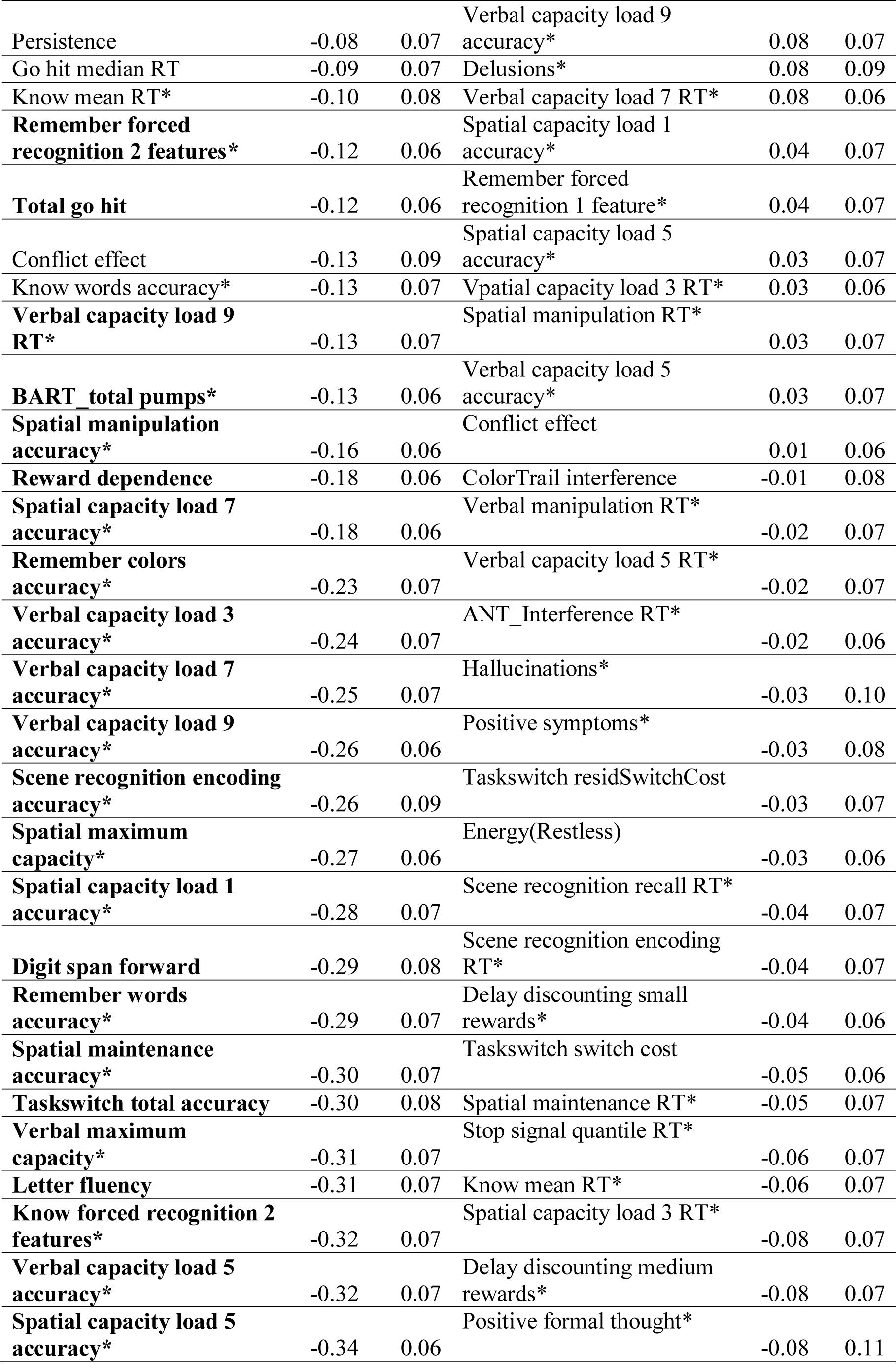
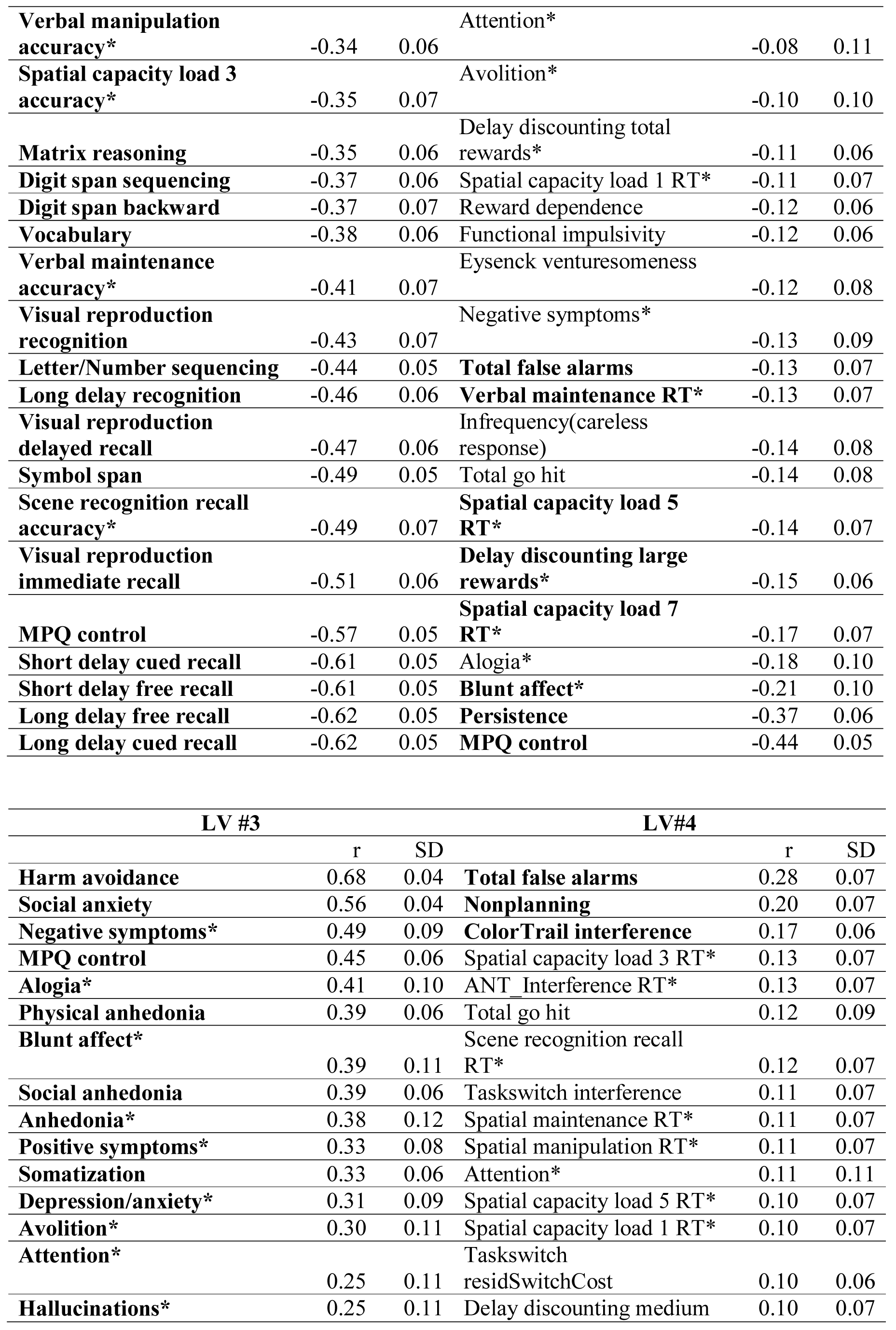
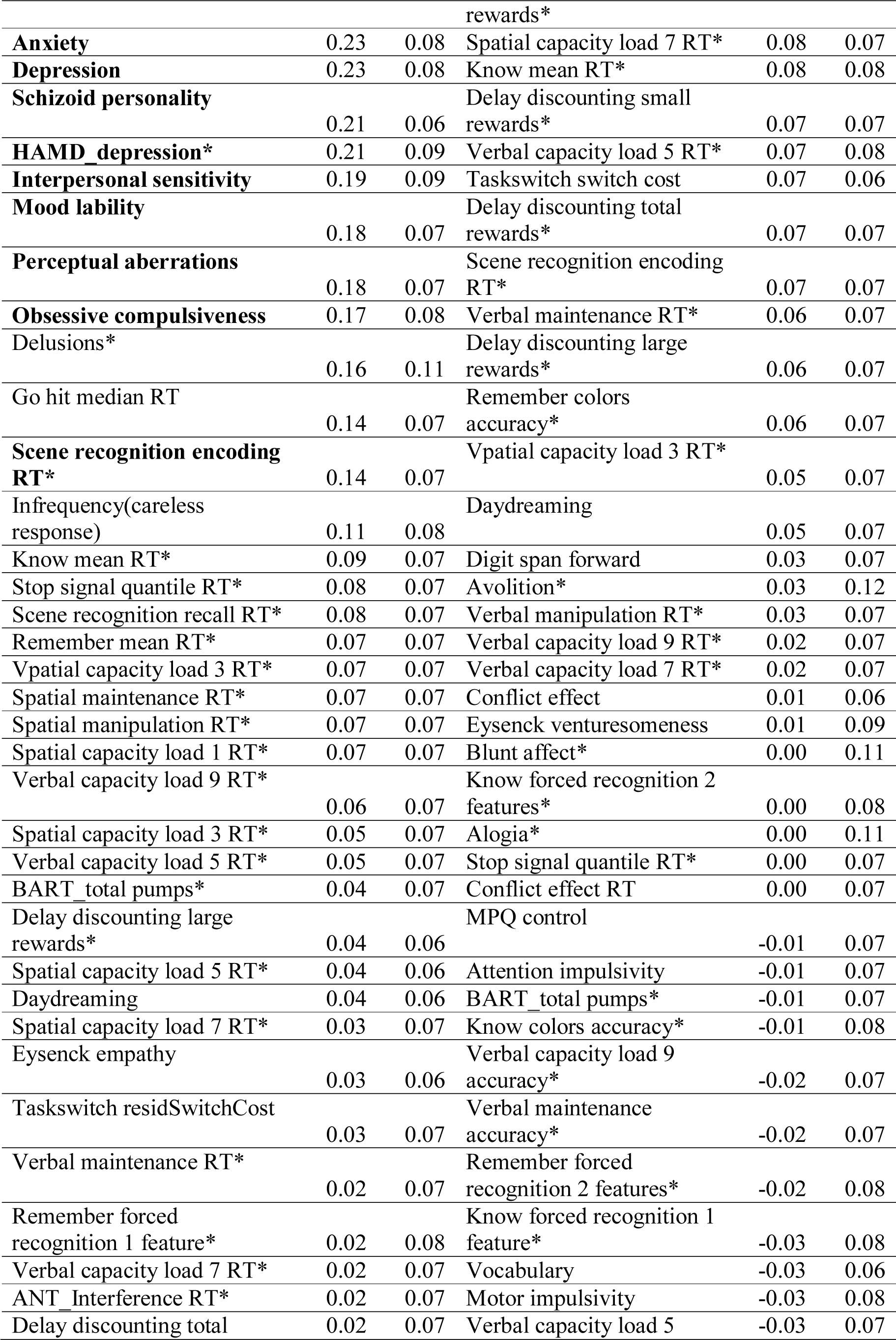
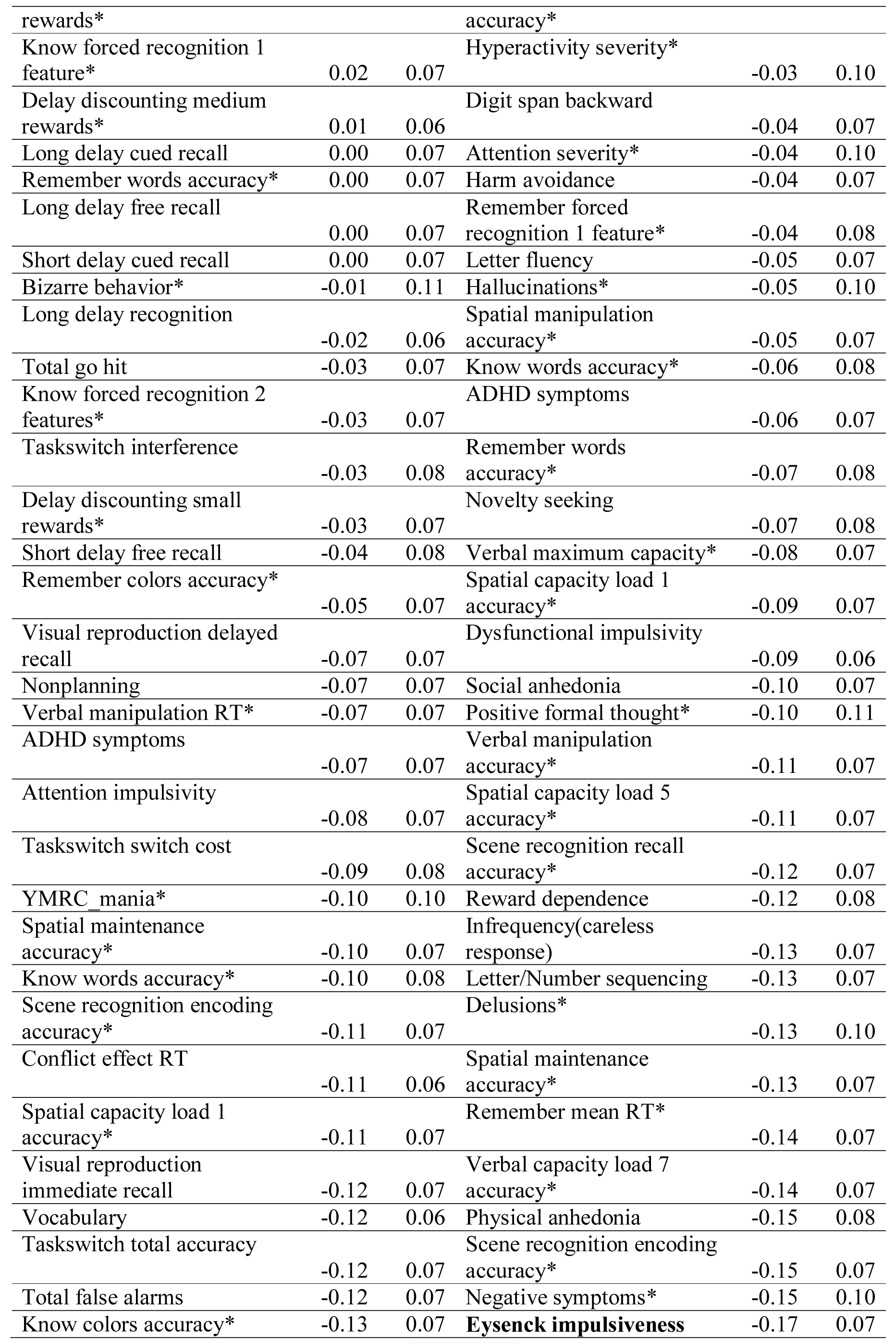
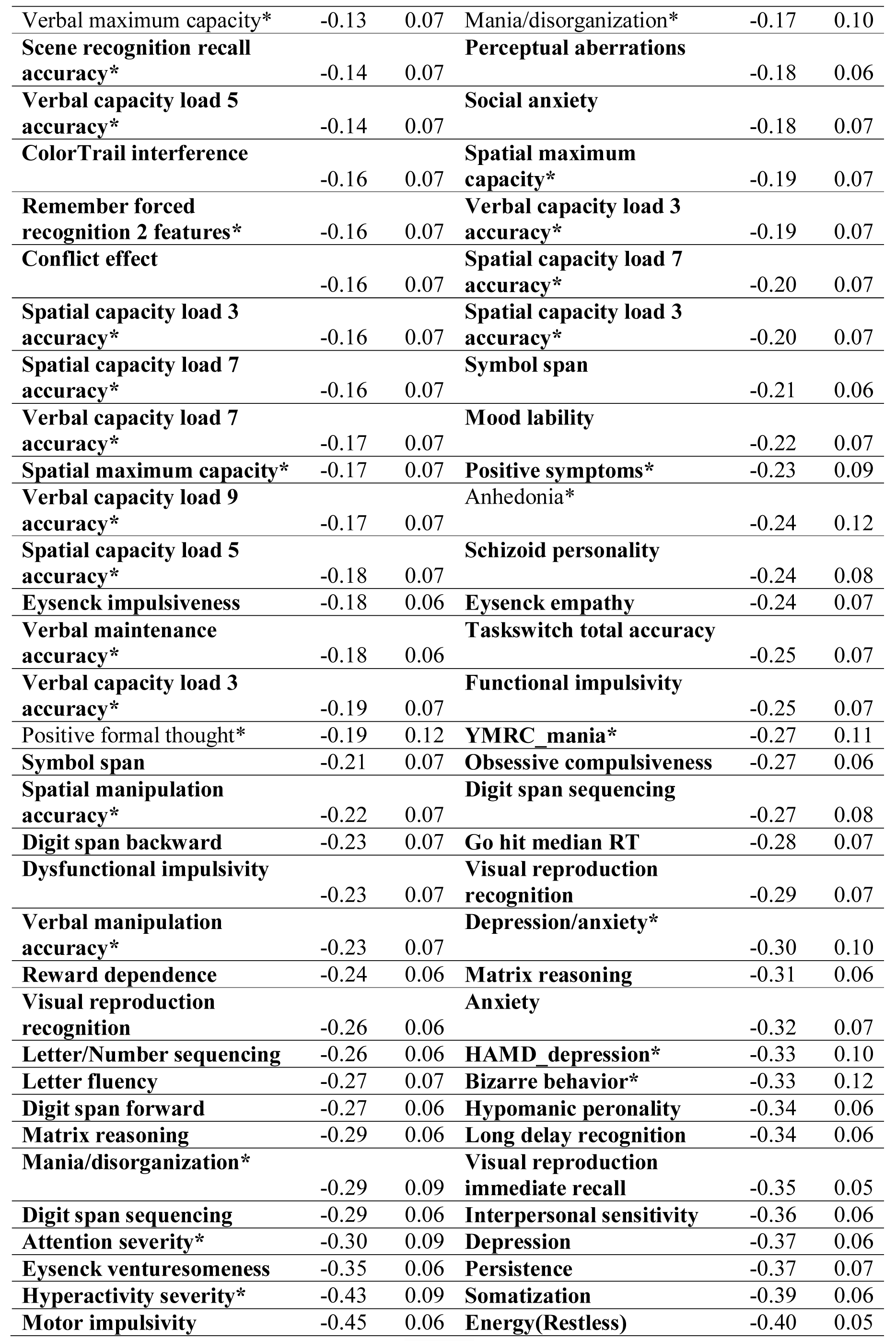
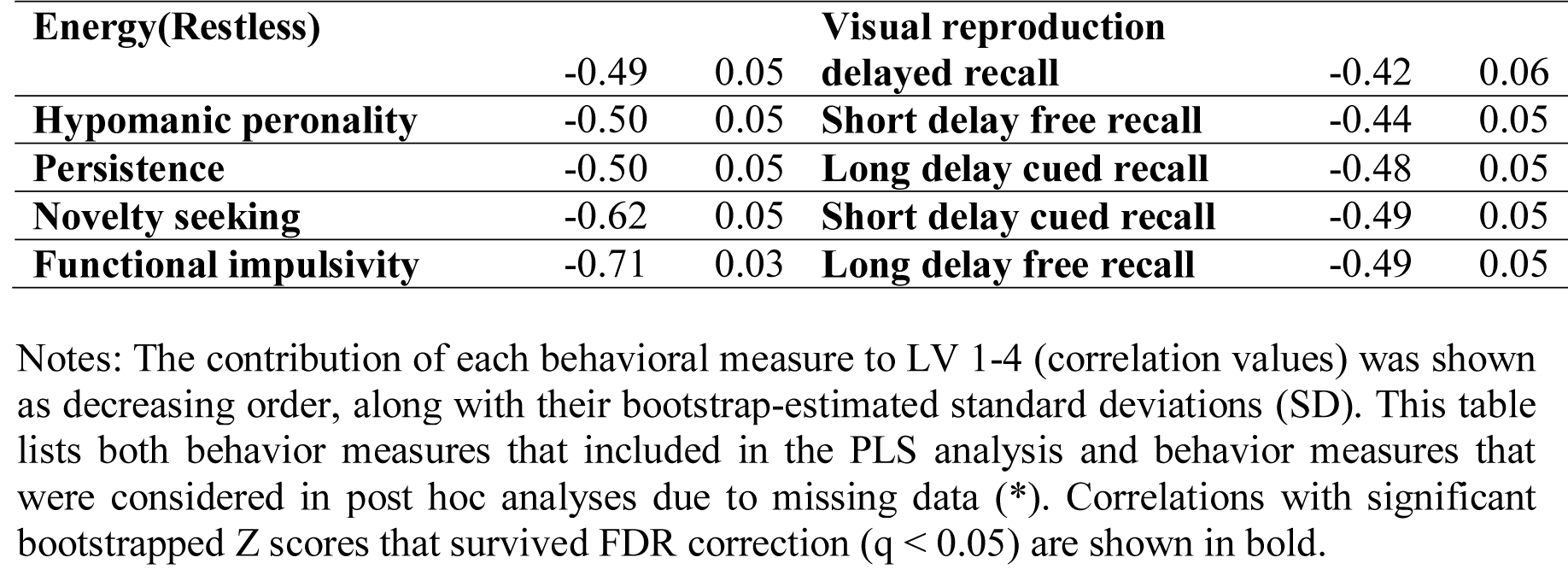
Correlations between subjects’ behavioral measures and their behavioral composite scores

